# Sensorimotor learning compensates for irreducible perceptual bias

**DOI:** 10.64898/2026.06.16.732722

**Authors:** Cléo L Schoeffel, Guilhem Ibos, Anna Montagnini, Guillaume S Masson

## Abstract

Because of the uncertainties present in any sensory inflows, we experience perceptual biases that often contaminate sensorimotor transformations. How these biases can be prevented through perceptual learning or adaptation is an open question in inference theories of perception and motor control. We revisited this debate using a classic example of low-level perceptual bias, the aperture problem in motion perception and its consequences for smooth pursuit. In a visuomotor tracking paradigm, participants pursued left-, right-tilted, or upright Gabor patches moving horizontally. As expected, initial pursuit direction was biased toward the oblique directions. This initial pursuit error remained unchanged when participants repeatedly tracked a single tilted Gabor during training sessions conducted over several days. By contrast, post-training error correction became faster for the trained Gabor orientation, but not for its untrained, mirror-symmetric counterpart. This change in dynamics was explained by the emergence of a delayed compensatory pursuit response, best revealed when testing with an upright Gabor, a stimulus that would normally elicit unbiased tracking. This corrective eye movement was selective for both shape and motion features. It lagged by ∼40 ms after pursuit onset, suggesting that it was triggered by the biased motor command itself rather than by visual reafference. These results support a dissociation between low-level sensory computations, which appear resistant to modification over the timescale of our training protocol, and internal models of sensorimotor transformation, which can be rapidly updated to compensate for irreducible but predictable sensory-driven perceptual biases.

## INTRODUCTION

Our perceptual systems are prone to systematic biases and illusions because sensory inputs are inherently uncertain ^1,2^. This question bears directly on perception as Bayesian inference where sensory likelihoods are combined with prior knowledge ^1,3^. Such uncertainties and biases are strong enough to influence our actions ^4,5^. A longstanding debate is whether these biases reflect fixed properties of sensory processing, such as hardwired prior expectations in the brain’s inference machinery, or whether they can be rapidly modified through experience. Yet evidence that perceptual learning reduces such biases remains sparse and controversial ^6–10^.

A striking illustration of this problem is the aperture problem: when a moving elongated contour is viewed through a limited receptive field, only the component of motion orthogonal to the contour’s orientation can be reliably estimated, leaving the true two-dimensional (2D) direction of motion undetermined. Such ambiguity results in a strong and persistent perceptual bias towards the contour-orthogonal direction ^11,12^. Moreover, any visual system resolves this ambiguity by integrating over space and time several local one-dimensional motion (1D) signals with different orientations ^12^ and unambiguous 2D cues ^13–15^. This integration process is well captured by a recurrent Bayesian inference framework in which initial motion estimates favor slow velocities, the so-called slow-speed prior ^16,17^, and then accumulate other 1D and 2D motion information to dynamically recover the true 2D global motion direction ^18,19^. Such dynamics are well represented at cellular level in the properties of many motion direction cells in macaque area MT ^14,20,21^ and pervade the ocular tracking system ^14,22–24^. In both macaques and humans, smooth pursuit of tilted moving 1D edges or objects is first initiated towards the direction orthogonal to the 1D edge, followed by a gradual correction over approximately 300 ms ^14,23,24^. Computational models of smooth pursuit further show that the dynamic resolution of the aperture problem in the oculomotor system can be understood as an optimal combination of sensory and predictive signals, implemented through internal models that are continuously updated by visual error signals ^25,26^. Overall, this example illustrates eye movements as simple but sensitive sensorimotor transformations that can be used to track the consequences of sensory uncertainties and their dynamical correction.

A long-standing and unresolved question is whether these motion perception biases can be reduced or eliminated through repeated sensory exposure. This could be seen as a form of perceptual learning broadly defined as experience-dependent improvement in sensory or sensorimotor processing ^8^. This question has profound theoretical implications: if a bias reflects a hardwired prior embedded in the architecture of sensory processing, it should be resistant to experience-dependent modification. Conversely, if priors are flexible statistical summaries updated by experience, repeated exposure to specific motion properties should reshape them. In the first case, sensorimotor systems must find other computational solutions to reduce any resulting behavioral biases while, in the latter, sensory updating would be sufficient. The existing literature is divided. A handful of perceptual studies reported that training reduces direction or speed perception biases, supporting the prior updating ^6,8,9^, while others claimed the opposite ^7,10^. Critically, this debate has been driven mostly by perceptual studies using discrete perceptual reports, which cannot dissociate the initial sensory bias from its subsequent correction, and conflate sensory and decisional contributions to behavior ^10^.

Here we reappraise this controversy using smooth pursuit eye movements. Unlike perceptual reports, smooth pursuit gives quantitative access to the initial direction error, reflecting the strength of the slow-speed prior at pursuit onset, as well as to the within-trial correction dynamics. We can thus decipher whether learning modifies the bias itself, the speed of its correction, or both ^14,24^. Inspired by previous perceptual studies ^9^, we designed a perceptual learning-like (**Fig.1)** paradigm in which participants repeatedly pursued a Gabor patch of a single 1D orientation moving either right or leftward at constant speed. We measured the dynamics of pursuit before and after training using Gabor patches of several orientations to disentangle learned and unlearned visual features. Participants ran several hundred trials, and the task was repeated over several sessions to probe learning over trials, blocks and days. We report that the initial direction error remains unaffected by training. However, the oculomotor system learns a predictive compensatory motor command, specific to the trained orientation and direction, that accelerates error correction across sessions. These results suggest a dissociation between the low-level sensory Priors that cannot be modified on the timescale of our experimental training sessions, and internal models of sensorimotor transformation that can be quickly updated to compensate for sensory-driven biases.

**Figure 1.**
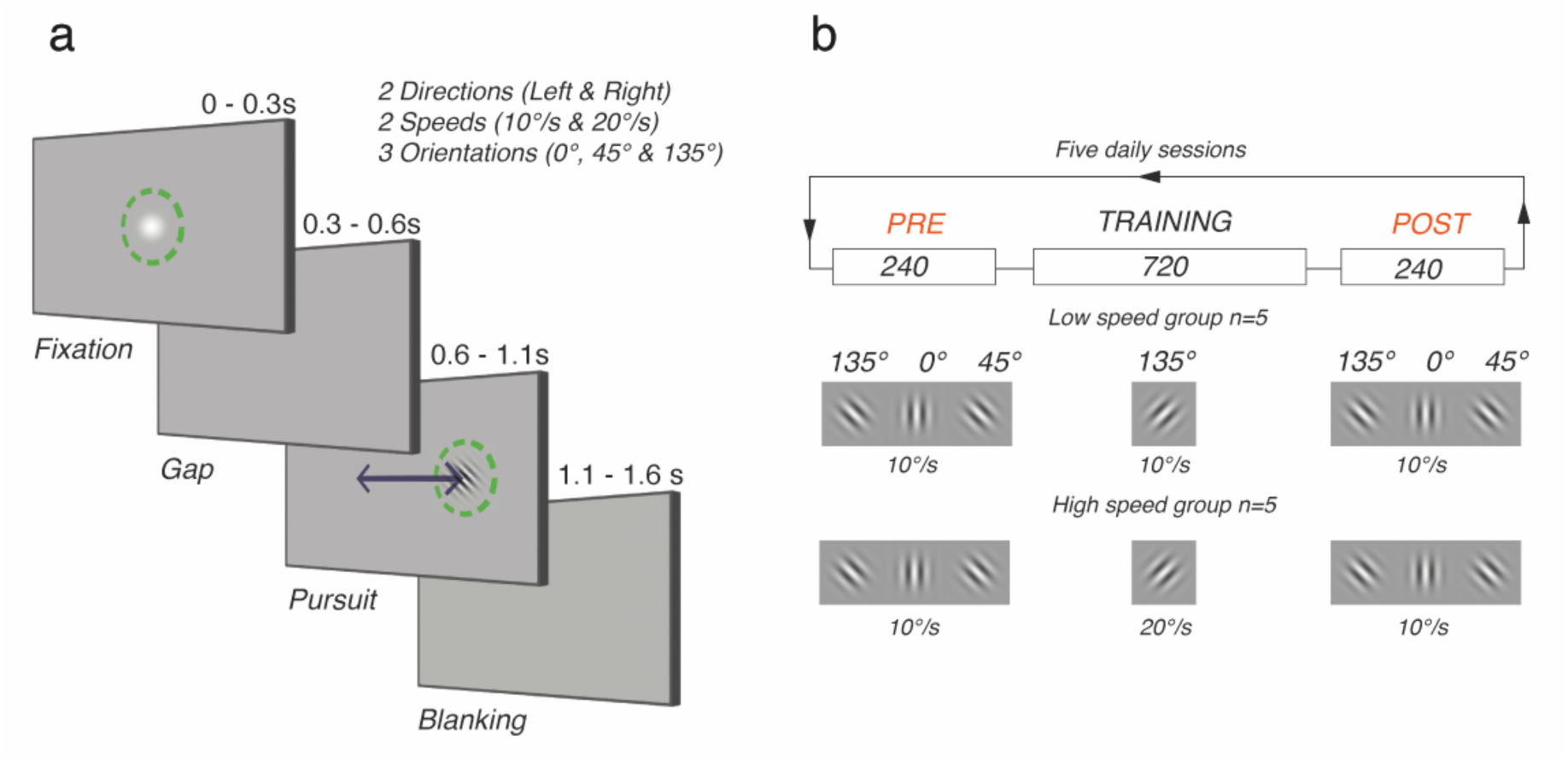
Experimental paradigm and training protocol. (**a**) Trial structure. Each trial began with a central fixation period (0–0.3 s) during which participants fixated a Gaussian blob. Following a gap period (0.3–0.6 s) in which the fixation stimulus disappeared, a Gabor target appeared at the centre of the screen and began translating horizontally (0.6–1.1 s). Participants were instructed to track the target as smoothly and accurately as possible. The target was then extinguished (blanking, 1.1–1.6 s) and the fixation spot reappeared, signalling the start of the next trial. The green dashed circle indicates the fixation window used to monitor gaze position online and was not visible to participants. (**b**) Training protocol. Each session consisted of a PRE block (240 trials), a TRAINING block (720 trials), and a POST block (240 trials). In PRE and POST blocks, all three orientations (135°, 0°, 45°) were fully interleaved and presented at 10°/s. During the TRAINING block, only the 135° orientation was presented. Two groups of five participants each underwent training at different speeds: the low speed group (n=5) trained at 10°/s, and the high-speed group (n=5) trained at 20°/s. Both groups completed 5 sessions of this protocol.

## RESULTS

We designed a learning paradigm to test how the aperture problem biases tracking performance and whether this bias changes after learning a specific orientation–direction association (**Fig. 1**). The paradigm was inspired by previous work claiming that biases in perceived motion direction can be eliminated through perceptual learning ^9^. Participants were asked to pursue with their eyes a Gabor target moving either to the right or to the left for 500 ms (**Fig. 1a**). During a pre-learning baseline session, three target orientations were randomly interleaved: 45°, 0°, and 135° counterclockwise. **Fig. 2a** illustrates, for one participant, the mean horizontal (left) and vertical (right) eye-velocity profiles for each target orientation and motion direction. With tilted (45° or 135°) targets, pursuit was initiated along both the horizontal and vertical axes, corresponding to an oblique direction roughly orthogonal to the target orientation. For instance, a 45° target moving to the right (solid pink lines) triggered an initial tracking response directed rightward and upward, as predicted by the aperture problem. Responses to leftward and rightward motion were mirror-symmetric, with the sign of the vertical component reversing with direction for a given tilt. By contrast, a vertical (0°) target moving in the same direction elicited a purely horizontal ocular response, with no vertical component. With tilted targets, vertical eye velocity peaked at around 200 ms after target motion onset before gradually returning to zero. As expected, horizontal eye velocity was slower for tilted Gabors than for the vertical one. Each of the five experimental sessions consisted of three successive blocks: a PRE-training block (240 trials, all three orientations interleaved), a training block (720 trials, 135° orientation only, presented repeatedly), and a POST-training block (240 trials, all three orientations interleaved). This structure allowed us to measure the aperture-induced bias and its correction before and after each training block, and to track how these measures evolved across the five sessions (**Fig. 1b**; see also Methods).

**Figure 2.**
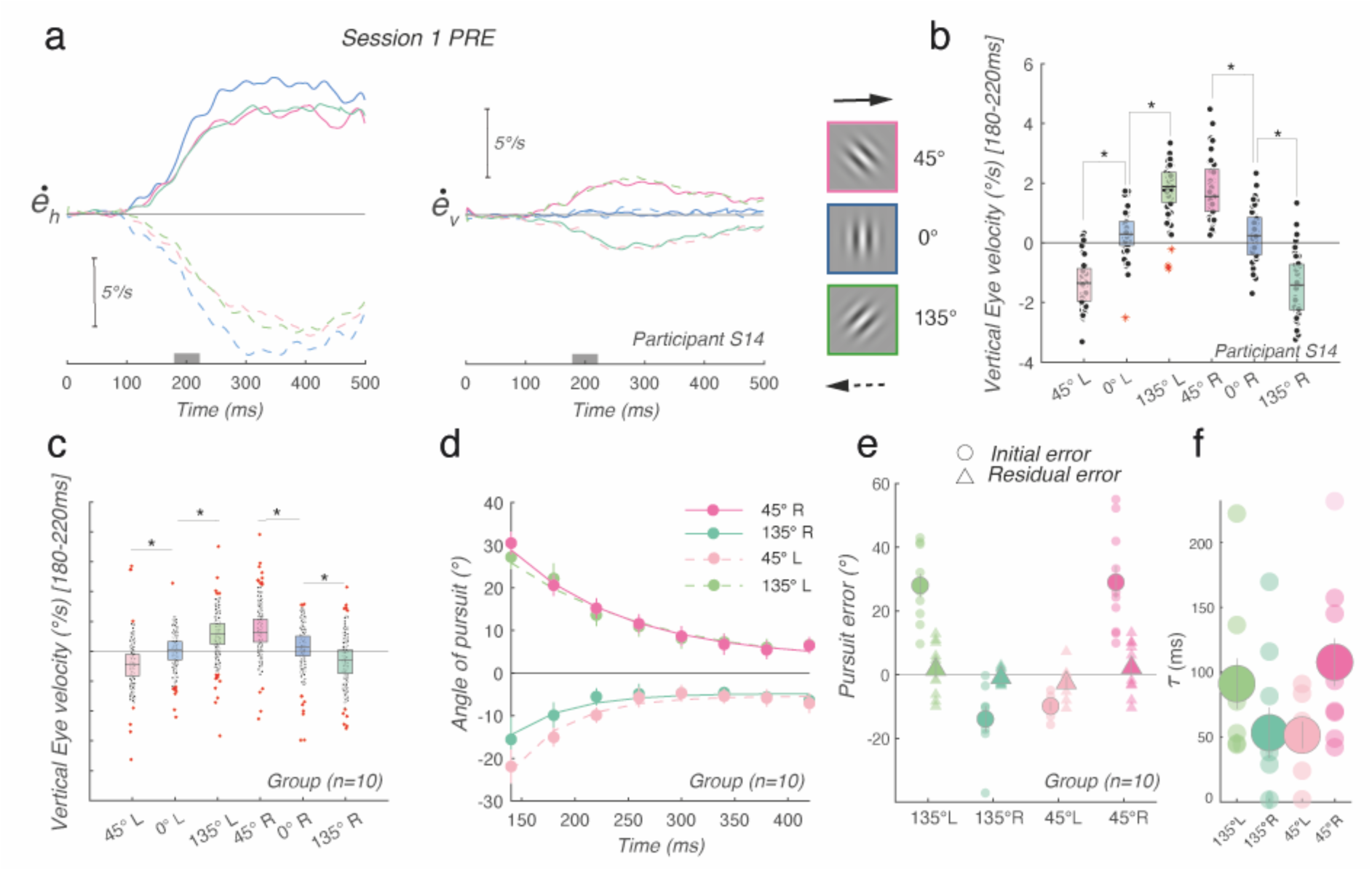
Aperture problem and smooth pursuit: initiation bias and correction dynamics. **(a)** Mean horizontal (left) and vertical (right) eye velocity profiles for one representative participant (*Participant 14*), Session 1 PRE, for all three Gabor orientations (blue: 0°, pink: 45°, green: 135°) and both motion directions (rightward: solid, leftward: dashed). **(b)** Mean vertical eye velocity in the [180,220] msec window for the same participant and session, shown for all orientation-direction combinations. * indicates p<.001 (Holm corrected) **(c)** Mean vertical eye velocity in the [180,220] msec window for all participants shown for all orientation-direction combinations for the PRE-training block of Session 1. * indicates p<.001 (Holm corrected) **(d)** Exponential fits to the pursuit angle time course ([120,440] msec) for the group-averaged pursuit angle time course (n=10), for oblique orientations and both directions (rightward: solid, leftward: dashed). R² > 0.93 for all fits. **(e)** Exponential fitting parameters: initial error (*a*, circles) and residual error (*c*, triangles) for the 135 and 45° tilt conditions. **(f)** Time constant (τ) for all subjects and all tilted Gabor conditions. Error bars denote ± SEM.

### The aperture problem impacts pursuit direction

Smooth pursuit unfolds in two phases: an initial open-loop phase, driven purely by visual motion signals before sensory feedback can influence the response, and a subsequent closed-loop phase during which the system detects and corrects the velocity (direction and speed) error. The initial phase reflects the output of the low-level motion processing. First, to quantify the initial pursuit bias, we computed the mean eye velocity over the (180-220) msec time window. While vertical eye velocity was near zero for vertical targets, it was significantly modulated for tilted orientations in a direction-dependent manner, as shown for one, illustrative naive participant (*Participant S14*, **Fig. 2b**). Since all participants completed identical pre-training blocks prior to any learning manipulation, data from all 10 participants were pooled for this baseline characterization regardless of their subsequent training group assignment (**Fig.2c**). A two-way repeated-measures ANOVA (n=10) revealed no significant main effect of orientation (F(2,18) = 1.95, p= .17, BF_10_ = 0.038) or direction (F(1,9) = 3.92, p = .079, BF_10_ = 2.59) when considered independently, but a highly significant Orientation x Direction interaction (F(2,18) = 55.62, p<.001, BF_10_ = 8.04×10^23^), confirming that the initial pursuit bias depended jointly on the target’s orientation and motion direction. Post-hoc paired t-tests (Holm corrected) showed that all three orientations differed significantly within each direction (all p<.005), while mirror-symmetric conditions (135° leftward versus 45° rightward, 135° rightward versus 45° leftward, and 0° leftward versus 0° rightward) did not significantly differ after correction (all p>.06), confirming the mirror symmetry of the direction bias.

Next, we measured how direction errors evolved over time by computing pursuit direction angle in successive 40 ms time bins, ranging from 120 to 440 msec after motion onset. **Fig.2d** plots these temporal dynamics for all participants, for all tilted conditions. The pursuit direction error decayed progressively towards a small, nearly zero asymptote ^27^, reflecting the well-known within-trial correction of the aperture problem. This decay was well captured by an exponential function of the form:

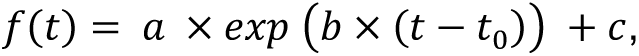

with R² values ranging from 0.88 to 0.97. Parameters *a* and *c* render the initial and final tracking error, respectively, and 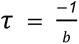 is the decay time constant. At the group level (**Fig 2d**), exponential fits were obtained for n=7-10 participants per condition after exclusion of invalid fits. The mean best-fit parameters were for the initial error amplitude *a* = 21.4±14.3°, the time constant *τ* = 77.5±56.6 msec, and the residual error *c* = 5.6±3.8°. The initial error *a* (circle in **Fig 2e**) showed a highly significant Orientation x Direction interaction (F(1,5) = 66.25, p<.001, BF_10_=1.43×10^13^), with all pairwise comparisons surviving Holm correction (all p<.005), confirming that the amplitude of the initial bias depends strongly on the specific orientation-direction combination. The time constant (τ) was log-transformed due to its skewed distribution. These log-transformed values also showed a significant interaction (F(1,5)=12.18, p=.017, BF_10_=5.51×10^5^), indicating that direction error dynamics differ across conditions, although individual post-hoc contrasts did not survive correction **(Fig.2f)**. Crucially, the residual error *c* (triangles in **Fig 2e**) showed no significant effect of orientation (*F*(1,6) = 0.38, *p* = .56, BF_10_ = 0.21), direction (*F*(1,6) = 3.32, *p* = .12, BF_10_ = 0.43), or their interaction (*F*(1,6) = 0.88, *p* = .38, BF_10_ = 1.29), demonstrating that regardless of target orientation, participants ultimately corrected their tracking to near-zero directional error within each trial. This initial tracking error and its reduction set the stage for the key question of this study: whether perceptual learning can reduce the initial direction error induced by the aperture problem, and/or modify the speed or residual amplitude of its subsequent correction. Since motion direction had no significant effect on vertical eye velocity (as shown by the non-significant main effect of Direction and the mirror symmetry of left and right responses, see **Fig 2c-f**), leftward trials were sign-flipped and pooled with rightward trials for all subsequent analyses, effectively doubling the number of trials per condition.

### Training speed does not influence the learning outcome

Sotiropoulos et al. (2011) showed that training with faster speeds can modify the slow-speed prior and reduce perceived direction biases. To test whether a similar manipulation would affect oculomotor learning, we compared participants trained at lower speed (10°/s, *n* = 5) and high speed (20°/s, *n* = 5) using linear mixed-effects models with Group as a between-subject factor and Session (S1–S5) and Block (PRE, POST) as within-subject factors, tested separately for mean eye velocity ([140,180] msec time window), initial tracking error (*a*), and time constant (τ). None of the models revealed a significant effect of Group, Session, Block, or any of their interactions (all *p* > .22). Bayesian model comparison further provided evidence supporting the absence of group differences (all BF_10_ ≤ 0.14, i.e., BF_01_ ≥ 7). Accordingly, data from both training speed conditions were pooled for all subsequent analyses, over a total group of 10 participants.

### Training does not change initial pursuit direction biases

We then compared PRE- and POST-training blocks – the 240-trial blocks of interleaved orientations recorded immediately before and after each training block – across the five successive sessions. We focused on two main indices. First, early vertical eye velocity reflects the direction bias at pursuit initiation. Second, the time constant (τ) with which this initial error is reduced reflects the temporal dynamics of direction-error correction. **Fig. 3a** illustrates vertical eye-velocity profiles for one representative participant (*Participant 14*) for both PRE and POST blocks (upper and lower panels) in Sessions 1 (left) and 5 (right). Initial tracking velocity appears largely unchanged between PRE and POST blocks, both in the first and last sessions. Similar observations were made in the three intermediate sessions. To quantify this effect, we extracted mean vertical eye velocity in the (140–180) ms time window. **Fig. 3b** plots initial eye velocity (single-trial and mean data) for the same illustrative participant across all tilted Gabor conditions and sessions (S1–S5). Comparisons between PRE and POST initial pursuit biases showed little, if any, difference in either the first or the fifth session. Indeed, a single-subject three-way factorial ANOVA (Block × Session × Orientation) revealed a strong main effect of Orientation (F(2,1160) = 444.07, p < .001, BF_10_ = 5.5110^5^) and no significant effect of Block, Session, or any interaction (all p > .10, all BF_10_ ≤ 0.11). **Fig. 3c** shows the mean values across all 10 participants. At the group level, a three-way repeated-measures ANOVA with Block (PRE, POST), Session (S1, S5), and Orientation (0°, 45°, 135°) as within-subject factors revealed a highly significant main effect of Orientation (F(2,18) = 96.84, p < .001), confirming robust and stable orientation tuning across all conditions. Critically, neither the main effect of Block (F(1,9) = 3.60, p = .09) nor that of Session (F(1,9) = 0.37, p = .558) reached significance, and none of the two-way or three-way interactions were significant (all p > .14). Bayesian paired tests provided moderate evidence supporting the absence of a training effect across blocks (BF_01_ = 5.42), while the session comparison was inconclusive (BF_01_ = 1.22). Post hoc comparisons confirmed that all three orientations differed significantly from one another (all p < .001), while this orientation tuning was independent of both Block and Session. Similar analyses across all five sessions yielded the same conclusion: training did not affect initial pursuit direction biases with tilted Gabors. Altogether, these results indicate that the initial vertical eye velocity—and thus the direction bias at pursuit onset—was not reduced by training, regardless of the number of sessions completed. The aperture-induced bias at tracking onset therefore appears resistant to perceptual learning.

**Figure 3.**
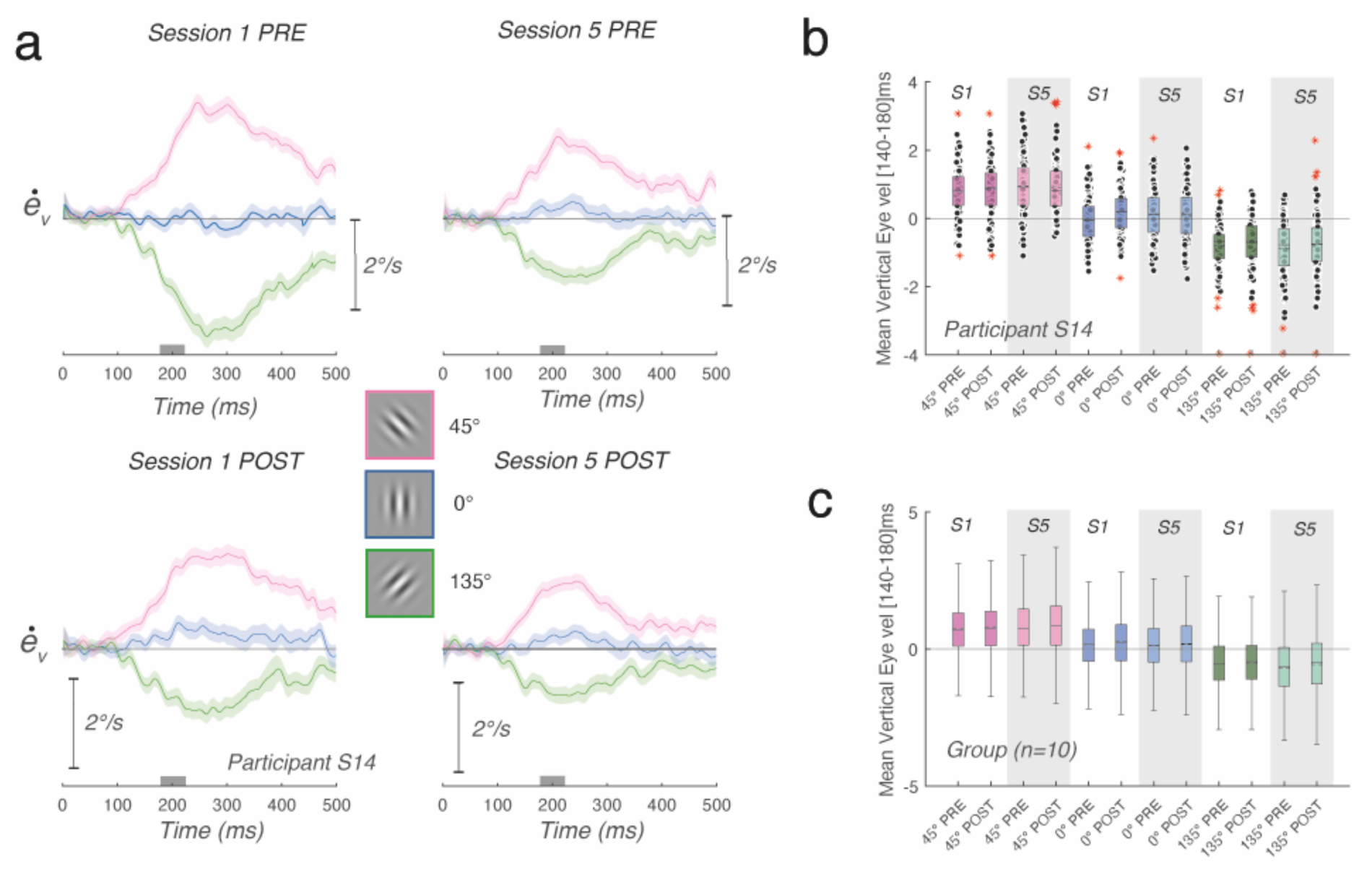
The initial direction bias at pursuit onset is not reduced by training. (**a**) Mean vertical eye velocity profiles for one representative participant (*Participant 14*, normal speed group) across all three Gabor orientations (blue: 0°, pink: 45°, green: 135°) with leftward trials sign-flipped and pooled with rightward trials (i.e., all data aligned to rightward motion), for PRE and POST blocks in Session 1 (top left and bottom left) and Session 5 (top right and bottom right). Shaded areas represent 85% confidence intervals. The grey box indicates the (140-180) ms analysis window. (**b**) Mean vertical eye velocity in the (140–180) ms window for Session 1 and Session 5, PRE and POST, for the same participant. (**c**) Same as (**b**) for all subjects (n=10). Colours and conventions as in Fig 2.

### Training speeds up pursuit direction correction

**Fig 4a** illustrates, for the same naive participant (*Participant 14*), the time course of the tracking direction error for the two oblique conditions (135°: green, 45°: pink), in the PRE (continuous lines) and POST (dashed lines) training blocks. Data from Session 1 and 5 are shown in dark and light shades respectively, and lines represent the best exponential decay fits to the data points. Compared to PRE blocks, POST blocks showed a faster exponential decay of the direction error, most clearly in Session 1. **Fig 4b** shows the group-level exponential fits to the pursuit angle time course (mean±SEM, n=10) for PRE and POST blocks of Session 1 and Session 5, for both oblique orientations. The progressive reduction of the time constant (τ) across sessions, confirmed by the statistical analyses below, is most apparent when comparing the fits between Session 1 and 5 for the learned orientation (135°). From individual exponential fits, we first extracted the initial tracking error parameter (*a*). Across all sessions and participants, *a* was strongly modulated by stimulus orientation (F(1,173)=66.7, p<.001, BF_10_ = 1.19×10^52^), with 135° and 45° conditions producing opposite-sign errors of comparable magnitude (mean ± SD : −14.37±8.7° and 15.97±8.7°, respectively), confirming the mirror-symmetric structure of the aperture-induced bias. Inspection of individual fits suggested a tendency for *a* to decrease across sessions, consistent with a gradual reduction of the initial error over the course of learning. However, neither the main effect of Block (PRE vs POST: F(1,173) = 0.59, p = .442, BF_10_ = 1.63×10^-11^) nor Session (F(4,173)=0.92, p=.456, BF_10_ = 7.79×10^-18^) reached significance, and none of the interactions were significant (all p>0.43, BF_10_ < 0.001). While the visual trend is suggestive, the initial amplitude of the direction error did not show statistically reliable change across the training protocol.

**Figure 4.**
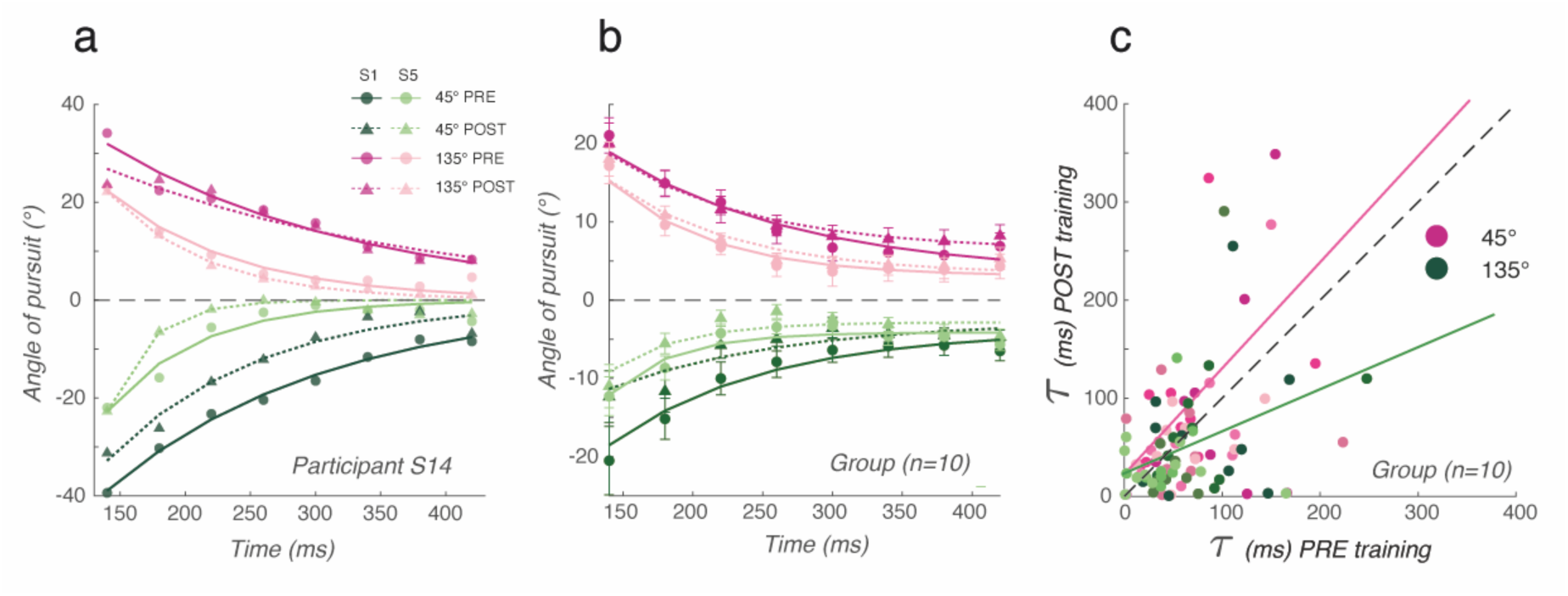
Learning accelerates direction error correction for the learned orientation across sessions. **(a)** Exponential fits to the pursuit angle time course for one representative participant (Participant 14), shown for Session 1 and 5, PRE (continuous lines) and POST (dashed lines) training blocks, for both oblique orientations (135°: green, 45°: pink). Dark and light shades correspond to Session 1 and 5 respectively. Filled circles and triangles: data points for PRE and POST respectively. Lines show best fits of the exponential decay function. **(b)** Exponential fits to the group-averaged pursuit angle time course (n=10), shown for PRE and POST blocks of Session 1 and 5, for the two oblique orientations. Conventions as in (a). **(c)** Correlation between PRE and POST (τ) values pooled across all subjects (n=10), shown separately for 45° (pink) and 135° (green). The dashed line indicates unity (PRE = POST). Points below the diagonal indicate smaller POST than PRE (τ) (Pearson correlations: 45°: r = 0.505, p < .001, R² = 0.26; 135°: r = 0.355, p = .014, R² = 0.13).

Overall, the time constant (τ) decreased both within and between the two illustrated Sessions. We ran the same analysis over 10 participants (**Fig.4b)**. Exponential fits confirmed a progressive reduction of the time needed to correct the pursuit error from Session 1 to Session 5, most clearly visible for the learned orientation (135°): (τ) decreased from 112-129 msec (PRE-POST range) in Session 1 to 46-50 msec in Session 5, compared to a more modest reduction from 103-137 to 74-91 msec for the unlearned 45° orientation. Given that (τ) values were not normally distributed, non-parametric tests were used throughout. A Friedman test revealed a significant effect of session (p<0.001), indicating that correction dynamics changed progressively over the course of the training protocol. *Post-hoc* comparisons showed that (τ) was significantly smaller in Sessions 4 and 5 compared to Sessions 1 and 2 (S1 vs S4: p=.016; S1 vs S5: p=.006, S2 vs S4: p=.016, S2 vs S5: p=.006), while no significant differences were found between adjacent sessions. By contrast, there was no significant effect of Block (Pre vs Post within session: Wilcoxon, p=.49, BF_01_ = 1.08). Likewise, the effect of Orientation (45° *vs* 135°) did not reach significance in the Wilcoxon test (p=.16), although Bayesian analysis provided moderate evidence for an orientation-related difference in *τ* (BF_10_ = 4.23). To validate these results using a model-free metric, we computed for each participant, session, block and orientation the time at which the direction error reached 50% of its initial value (t_50%_), a measure of correction speed that does not depend on assumptions about the shape of the decay function. A three-way repeated-measures ANOVA with Session (Session 1-5), Phase (PRE, POST) and Orientation (45°, 135°) as within-subject factors revealed a significant main effect of Session (F(4,36) = 2.74, p = 0.04), confirming that correction speed improved progressively across training sessions. The marginal means showed a clear reduction from Session 1 (160.7 ± 15.9 msec) to Session 5 (130.4±12.8 msec). Consistent with the (τ) analysis, neither the main effect of Phase (F(1,9) = 0.10, p = .76) nor Orientation (F(1,9) = 4.43, p = .06) reached significance, though the Orientation trend was in the expected direction (135°: 135.8±11.9 *vs* 45°: 159.1±8.1 msec). These model-free results confirm that the improvement in error correction dynamics across sessions is robust and does not depend on the specific assumptions of the exponential decay model.

To test whether training specifically modified correction dynamics for the learned orientation, we compared PRE and POST (τ) values across all participants. A constant relationship would indicate no systematic training effect, while a systematic shift below the unity diagonal would indicate training-induced reduction. Examination of the PRE *vs* POST (τ) correlation plot (**Fig 4c**) revealed a clear asymmetry between the two tested Gabor orientations. Both correlations were significant, indicating that individual correction dynamics were reliably preserved across blocks. However, the slope of this relationship differed markedly between the two orientations. For the unlearned orientation (45°, pink line), PRE and POST *τ* values clustered around the unity diagonal (r^2^=0.505, p<.001, r²=0.26, BF₁₀ = 1001), indicating that the individual dynamics were highly stable across blocks with no systematic shift. For the learned orientation (135°, green line), the correlation was significantly weaker (r^2^=0.355, p=.014, r²=0.13, BF₁₀ = 22.3), and data points shifted below the diagonal, with PRE (τ) being longer than POST (τ) across all participants. The dramatic difference in Bayes factors between orientations (BF₁₀= 1001 and 22.3) quantifies this asymmetry: learning partially disrupted the stability of individual correction dynamics specifically for the learned orientation, consistent with a genuine learning-induced acceleration of error correction speed.

To further characterize the dynamics of learning within the individual training blocks (**Fig. 5**), we compared mean pursuit direction between the first 20 trials (0-20) and late 20 trials (640-660) of each training session. For each selected trial, we computed both initial ([180,220] msec) and steady-state ([400,440] msec) time windows. Pursuit direction values were normally distributed in all conditions (Lilliefors tests, all p > .07). In Session 1, a 2×2 repeated-measures ANOVA with Phase (Initiation, Late) and Bin (First, Last) as within-subject factors revealed a significant main effect of Phase (F(1,9) = 15.77, p = .003), reflecting the established difference between initiation and late pursuit direction. The main effect of Bin (F(1,9) = 1.19, p = .30) and the Phase × Bin interaction (F(1,9) = 0.54, p = .48) were not significant. Planned paired comparisons, however, showed a significant first-to-last shift in late pursuit direction (Late phase: First = −7.31°, Last = −4.74°; t(9) = 2.60, p = .029, BF₁₀ = 2.70), but not in initiation pursuit (Initiation phase: First = −14.36°, Last = −14.24°; t(9) = −0.05, p = .96, BF₀₁ = 3.24), showing that initiation pursuit was left unchanged across the entire training block. This dissociation suggests that the faster correction develops selectively during the training block, resulting in lower final direction errors.

**Figure 5.**
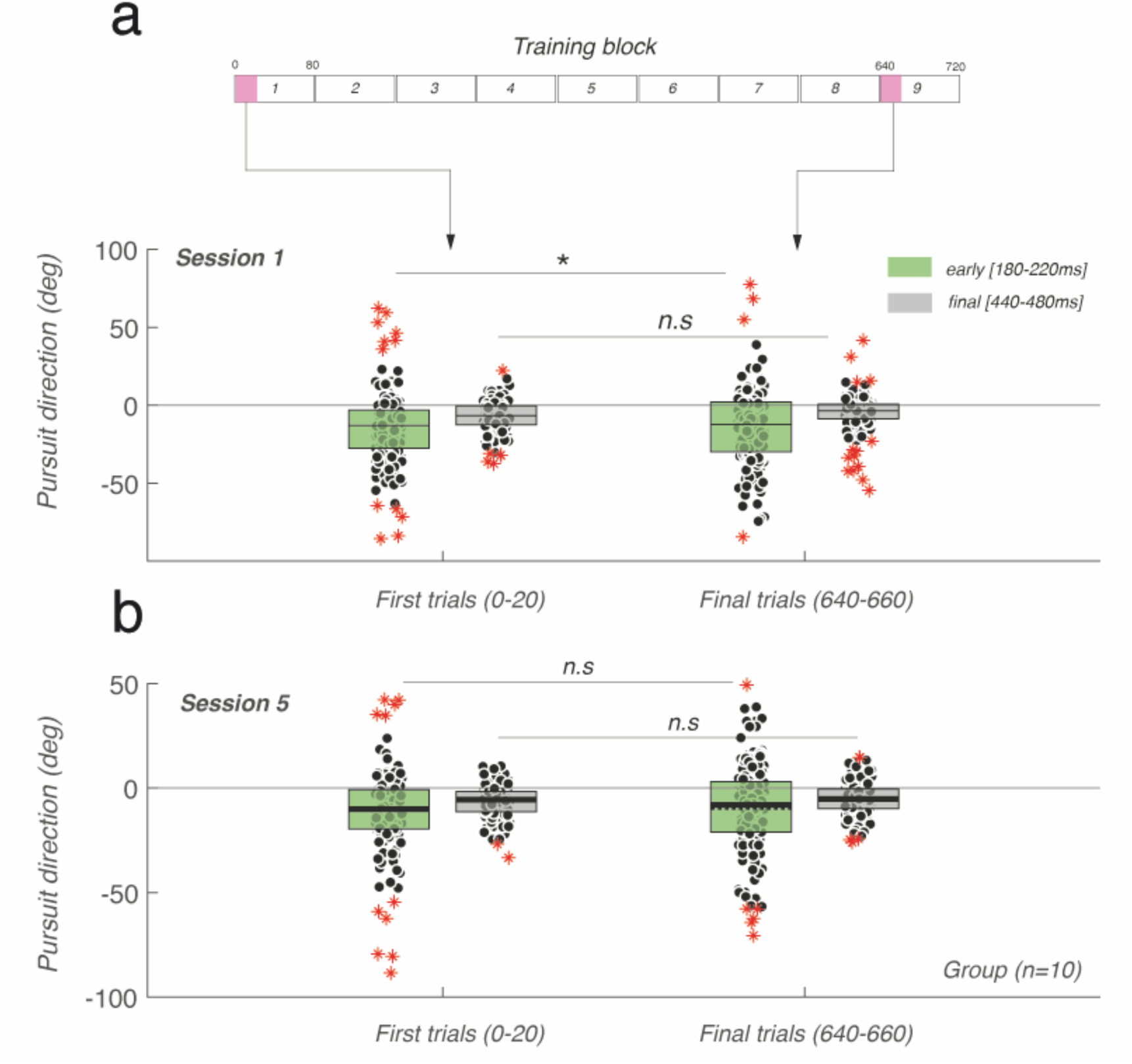
Within-session dynamics of pursuit initiation and residual error during training. **(a)** Pursuit direction at two different time windows ([180,220] msec) in green, [440,480] msec) in grey) during the first 20 trials (left) or late trials (right). The ruler illustrates the trial schedule for the training blocks (total 720 trials) during the first session. (**b**) Same plots, for the last training session (Session 5). Data are single trials (n=200) and mean pursuit direction across 10 participants.

In Session 5 (lower plot), the same analysis revealed a significant main effect of Phase (F(1,9) = 7.79, p = .021) but no main effect of Bin (F(1,9) = 1.06, p = .33) and no Phase × Bin interaction (F(1,9) = 0.14, p = .71). Neither the late nor the initiation window showed a significant first-to-last shift (Late: t(9) = −1.10, p = .30, BF₀₁ = 1.98; Init: t(9) = −0.79, p = .45, BF₀₁ = 2.49), and pursuit direction variability decreased markedly (Late Last SD: 1.76° in Session 5 *vs* 3.68° in Session 1). Together, these results indicate that the faster correction, which develops within Session 1, is already consolidated and stable from the first trials of Session 5, leaving little room for further within-session shifts.

### Training builds up a compensatory pursuit component

The results presented above show that the initial direction bias was unaffected by training, whereas the speed of tracking-error correction gradually improved across sessions, selectively for the trained orientation. We next asked whether this learning effect was truly stimulus-specific, or whether it reflected a more general compensatory response that could also be revealed in conditions for which the aperture problem itself predicts no directional bias. To address this question, we examined vertical eye-velocity profiles for the vertical Gabor (0°). As shown for the first PRE-training session, a vertical Gabor did not produce any vertical eye-velocity component (**Fig 2a**), because for a vertically oriented carrier the aperture-induced bias is aligned with the true 2D motion direction. Pursuit under these conditions is therefore strictly horizontal, as expected. However, as visible in **Fig 3a** for the same participant, after the first training session (POST, Session 1) a small but significant vertical ocular response was elicited by the vertical Gabor moving horizontally (blue curve). This vertical response was direction- and orientation-selective: it was opposite in sign to the pursuit bias elicited by the trained 135° target (**Fig. 6**, blue curves). Moreover, these post-training responses were delayed relative to the aperture-induced responses elicited by tilted Gabors (140 ± 23 *vs* 97 ± 12 msec) and peaked at ∼200 msec after motion onset. This compensatory response was already present before training in Session 1 POST and was further enhanced after training. Despite some idiosyncratic differences in amplitude, it was observed in all participants and in all post-training blocks. Across participants, vertical responses elicited by the 0° Gabor were upward for rightward motion (after sign-flipping and pooling leftward trials), that is, in the opposite direction to the downward vertical bias elicited by the trained 135° Gabor (**Fig. 6**).

**Figure 6.**
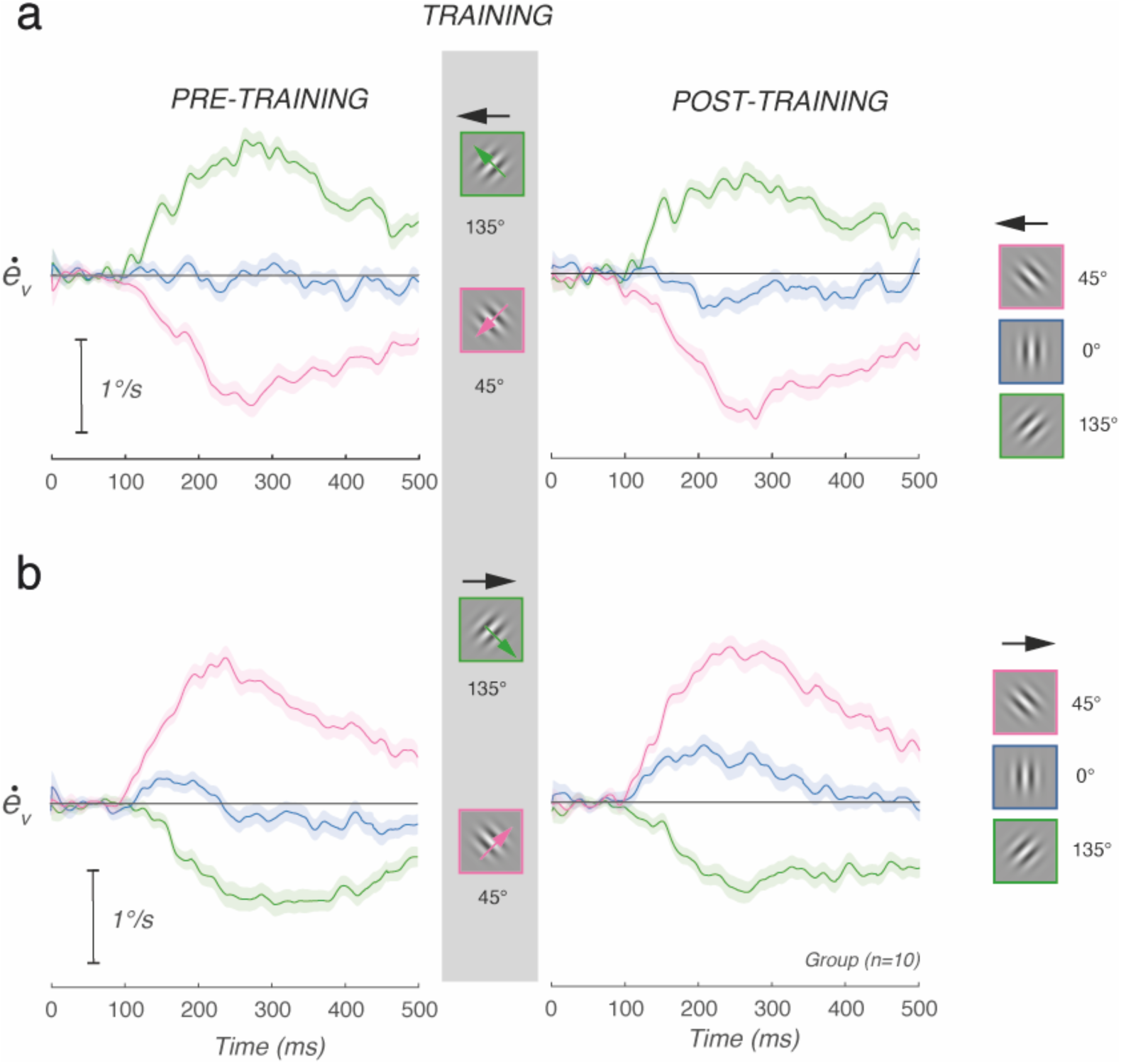
Mirror symmetry of the aperture-induced bias and learned compensatory response across motion directions. Mean vertical eye velocity profiles for all participants (n=10) separately for leftward **(a)** and rightward **(b)** motion directions, averaged across Sessions 1 PRE (left) and POST (right) blocks. Colours indicate Gabor orientation: blue: 0°, pink: 45°, green: 135°. Shaded areas represent 85% confidence intervals.

**Fig 7** further details these compensatory responses. **Fig 7a** plots mean vertical eye velocity profiles for each of the 5 Sessions (dark to light blue) and PRE and POST blocks (left and right-hand plots). In this illustrative, naïve participant (*Participant 14*), no vertical eye movement can be seen in the PRE block of Session 1, where vertical eye velocity remained near zero as expected (dark blue curve). By contrast, a small but consistent vertical eye velocity component emerged robustly in the first POST block and persisted across the 4 subsequent sessions. Moreover, a smaller but consistent vertical response to a 0° target was visible throughout the PRE blocks of sessions 2 to 5. **Fig 7b** plots the mean vertical eye velocity profiles across 10 participants and all 5 sessions for both PRE (continuous lines) and POST (broken lines) training blocks, for the 3 target orientations. The vertical responses for a 0° Gabor were clearly visible and consistent (**Fig 7b,** n=10), indicating a consistent effect across participants. Since the 0° response was consistently upward, it reduced the downward bias observed with the trained +135° target but increased the upward bias generated by the untrained +45° target. Moreover, these mean velocity profiles show that these compensatory vertical responses were delayed. These late consequences can be observed by comparing the continuous and broken lines for the tilted Gabor conditions (pink and green). We measured vertical pursuit latencies from the individual mean profiles, for each condition and block (**Fig 7c**). With oblique Gabors (45° and 135°), vertical eye movements were initiated at 97±12 msec (mean ± SD, n=200). By comparison, with the vertical Gabor (0°), vertical eye movements were initiated at 140±23 msec (mean ± SD, n=72). Vertical latencies for vertical Gabors were significantly different from both 45° and 135° orientations (paired t-tests, both p < 0.001, Bonferroni-corrected α = 0.025). Bayesian analyses revealed strong evidence for these differences (BF₁₀ = 9.355 and 14.691, respectively).

**Figure 7.**
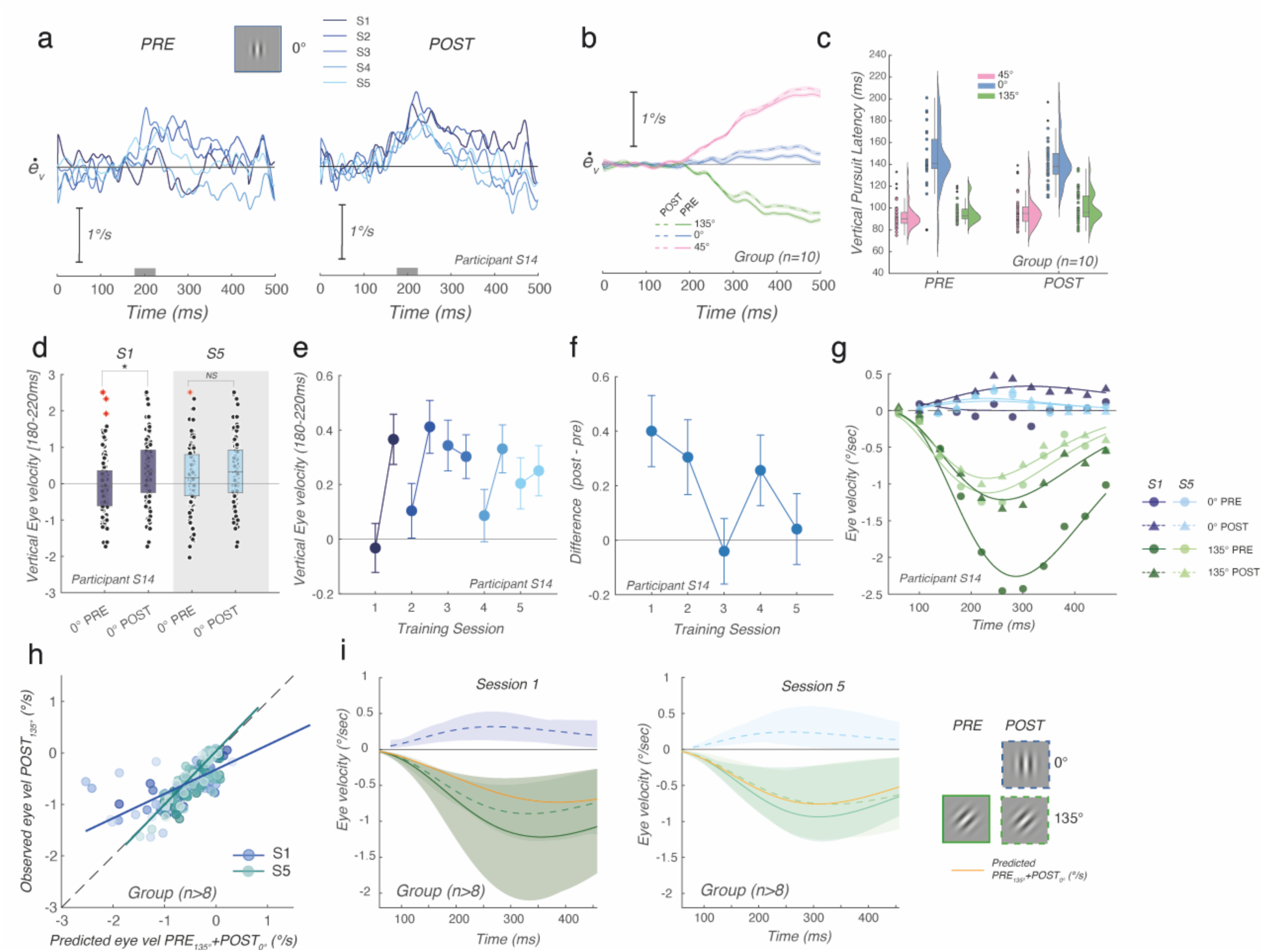
**Training induces a predictive compensatory motor response specific to the trained orientation.**(**a**) Mean vertical eye velocity profiles for the vertical Gabor (0°) only, for one representative participant (*Participant 14*), across all five sessions (S1: darkest blue, S5: lightest blue) for PRE (left) and POST (right) training blocks. (**b**) Mean vertical eye velocity profiles for all subjects and all sessions, for PRE and POST blocks, for 135°, 0°, and 45° conditions (PRE: solid, POST: dashed). (**c**) Vertical pursuit latency distributions for 0°, 45°, and 135° conditions, PRE and POST, across all subjects (n=10). (**d**) Mean vertical eye velocity in the [180,220] msec time window for 0°, Sessions 1 and 5, PRE and POST, for *Participant 14*. * indicates p<.01. (**e**) Mean vertical eye velocity ((180– 220) msec) across all five sessions for PRE and POST blocks (same participant). (**f**) Bootstrap estimates (1000 resamples with replacement) of the mean POST minus PRE difference in vertical eye velocity across sessions (same participant). (**g**) Best-fit Gamma functions to the vertical eye velocity time course for 0° and 135° conditions, Session 1 and Session 5, PRE and POST (same participant). (**h**) Correlation between predicted (PRE 135° + POST 0°) and observed (POST 135°) eye velocity per 40 msec time bin, for all subjects (n>8) pooled across sessions. Group-level Pearson correlations: S1: R²=0.668, p<.001, slope=0.47; S5: R²=0.823, p<.001, slope=0.911. **(i)** Temporal dynamics of observed and predicted vertical eye velocity for Session 1 (left) and Session 5 (right), averaged across subjects (n>8). Shaded areas represent 85% confidence intervals. The orange dashed line shows the linear prediction (PRE 135° + POST 0°). Inset illustrates the stimulus conditions.

We extracted the initial amplitude of vertical responses to a 0° target for each trial. Note that we used a later time window ([180,220] rather than the usual [140,180] msec) to quantify the 0° response, reflecting its longer onset. This window captures the 0° response at its peak expression. **Fig 7d** illustrates the mean vertical responses before (PRE) and after (POST) training, in both first and last session, respectively, for the same illustrative participant (*Participant 14*). A two-way factorial ANOVA (Session x Block) revealed a significant main effect of Block (F(1,376) = 9.29, p = .0025, BF₁₀ = 5.31), with POST vertical eye velocity being significantly greater than PRE (mean difference = −0.28°/s, 95% CI (−0.46, −0.10), p = .0023). Neither the main effect of Session (F(1,376) = 1.66, p = .198, BF₁₀ = 0.12) nor the Session x Block interaction (F(1,376) = 1.66, p = .198, BF₁₀ = 0.12) reached significance, with Bayesian analysis providing moderate evidence for the absence of both effects. Across 10 participants, the mean initial vertical response was significantly higher in POST than PRE blocks (main effect of Block: F(1,9) = 6.86, p=.028), and the magnitude of this PRE-POST difference depended significantly on session (Session x Block interaction: F(1,9) = 5.6, p=.042). Post-hoc comparisons revealed that the PRE-POST difference was significant in Session 1 (p=.009) but not Session 5 (p=.229), with mean velocities of 0.11°/s (PRE) vs 0.42°/s (POST) in Session 1, and 0.10°/s (PRE) vs 0.22°/s (POST) in Session 5 (**Fig 7d**). Bayesian analysis provided moderate evidence for PRE-POST difference (BF_10_ = 3.14), while supporting the absence of a main effect of session (BF_10_ = 0.42). To better illustrate the session-by-session dynamics underlying this group effect, **Fig 7e,f** show the mean initial vertical responses PRE and POST training for each session and their bootstrapped differences, respectively, for the same representative participant (*Participant 14*). This individual example is representative of the pattern observed across all participants. Over repetitive training, the PRE vertical responses built up while POST responses remained steady, further reflecting the adaptive nature of these compensatory responses.

To confirm that this compensatory response was not driven by a small subset of participants, we performed the same two-way ANOVA (Session × Block) independently for each of the 10 participants (Table 1). The Block (PRE vs POST) effect was individually significant in 7 of 10 participants, with extreme Bayesian evidence (BF₁₀ > 100) in 4 of 10. As a complementary, model-free test, we also compared single-trial vertical eye velocity between PRE and POST blocks (pooled across all 5 sessions) for each participant using a Wilcoxon rank-sum test. This confirmed the effect in the predicted direction in 8 of 10 participants, reaching individual significance in 8 of 10, with moderate-to-extreme Bayesian evidence (BF₁₀ > 3) in 6 of 10 (Table 1). One participant (S12) showed a small, non-significant effect in the opposite direction, and one participant (S03) showed a significant effect in the opposite direction. These single-subject analyses demonstrate that the compensatory response, while modest or absent for a minority of individuals, is a robust and highly prevalent phenomenon across our sample rather than being driven by a subset of participants.

**Table 1.** The compensatory 0° response is robust at the single-subject level. For each of the 10 participants, we tested whether vertical eye velocity to the 0° Gabor ([210,260] msec window, capturing the response at its peak expression) was greater in POST than PRE training blocks. Left columns: two-way ANOVA (Session × Block), the same design used for the representative participant in Fig 7d. Right columns: Wilcoxon rank-sum test on single-trial values pooled across all 5 sessions.

| Subject | ANOVA<br>F(Block) | ANOVA<br>p(Block) | ANOVA<br>BF10(Block) | Wilcoxon<br>z | p | BF10 | Cohen's d |
| --- | --- | --- | --- | --- | --- | --- | --- |
| S01 | 87.50 | 8.89e-20 | 6.12e+16 | 7.87 | 3.64e-15 | 3.95e+13 | 0.606 |
| S02 | 17.19 | 3.82e-05 | 209.37 | 4.21 | 2.52e-05 | 522.47 | 0.325 |
| S03 | 2.54 | 0.1114 | 0.14 | -2.28 | 0.0223 | 0.41 | -0.136 |
| S08 | 6.98 | 0.0084 | 1.21 | 2.72 | 0.0066 | 7.39 | 0.218 |
| S12 | 1.06 | 0.3041 | 0.07 | -1.30 | 0.1945 | 0.17 | -0.095 |
| S05 | 7.76 | 0.0055 | 1.99 | 3.08 | 0.0021 | 5.43 | 0.234 |
| S07 | 29.48 | 8.26e-08 | 9.21e+04 | 5.05 | 4.50e-07 | 7.30e+04 | 0.446 |
| S13 | 3.42 | 0.0649 | 0.22 | 2.48 | 0.0130 | 0.49 | 0.148 |
| S14 | 17.73 | 2.87e-05 | 262.25 | 4.23 | 2.34e-05 | 648.70 | 0.315 |
| S15 | 6.90 | 0.0088 | 1.32 | 2.69 | 0.0071 | 2.99 | 0.220 |

Finally, we evaluated the consequences of such vertical responses to the pursuit dynamics for tilted targets. As shown above, the vertical ocular responses to a 0° Gabor had latencies ∼40 msec longer than those for 45/135° Gabor. During this first 40 msec, responses are purely horizontal before deviating vertically, contrary to what was found with the tilted Gabors. To account for these differences, we changed our model. First, the first data point ([80,120] msec) was omitted for the 0° Gabor condition, for each block and session. Second, we used vertical eye velocity and not pursuit direction as an index of direction error, to capture the full dynamics of the vertical component. Third, we changed to a Gamma function, which allowed us to keep the same timeline across tilted and upright Gabor conditions while rendered the initial rise in vertical responses and their subsequent exponential decay. The fitted function was:

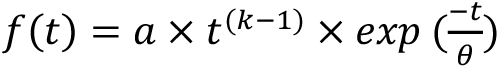

where *a* is a scaling factor, *k* the shape parameter controlling the rise time, and τ the scale parameter controlling the decay. Gamma function fits with r²< 0.5 were excluded. This criterion retained 75% of fits across all subjects, sessions, and conditions. r² values for retained fits ranged from 0.51-0.98 (mean ± SD: 0.82±0.13). **Fig 7g** plots for the same illustrative *Participant 14*, the time course of the vertical component velocity with either a vertical Gabor (blue curves) or a 135° Gabor (green curves) for both PRE and POST blocks of the first and last session. Continuous lines are the best-fit Gamma function (see Methods). One can see that, as vertical responses to the 0° target developed from Session 1 (POST) to 5 (PRE and POST), the vertical error to the 135° target reduced from PRE to POST training blocks, with this reduction being smaller with the last training session.

We reasoned that the POST training response to the learned orientation (135°) should equal the sum of the pre-learning aperture-induced bias (PRE 135°) and the newly learned compensatory response (POST 0°). We directly tested this hypothesis by computing, for each time bin, the predicted POST 135° response as PRE 135° + POST 0° and comparing it to the actually observed POST 135° response, in both Sessions 1 and 5. **Fig 7h** shows these relationships for each participant. We computed Pearson correlations between predicted and observed POST 135° eye velocities across all participants. The agreement between predicted and observed was striking: the two were strongly correlated in both Session 1 (r² = 0.668, p < .001, slope = 0.47) and Session 5 (r² = 0.823, p < .001, slope = 0.911), with the goodness of fit improving across sessions. The observed POST 135° response was systematically smaller than predicted, consistent with sublinear superposition, but the strong and increasing correlations confirm that the compensatory vertical response learned for the 0° Gabor is not merely a curiosity, but a quantitatively meaningful and predictive component of the improved correction dynamics observed at the trained orientation after learning.

To further confirm this relationship, we generated theoretical Gamma functions of temporal dynamics from the mean best-fit parameters of the PRE_135°_, POST_0°_ and POST_135°_ conditions. **Fig 7i** illustrates them for both the first (left-hand) and the last (right-hand) sessions. From these functions, we computed a predicted temporal dynamic of the POST 135° pursuit response (orange curves). In the first session (left plot), the comparison confirmed the sublinear interactions: the orange curve is lower than the dashed green one. After 5 training sessions, the temporal dynamics for the response to a 135° target can be fully explained by the effect of the aperture problem (sensory estimates) and the compensatory eye movements. The predicted temporal dynamics closely matched the observed POST_135°_ response in both sessions (Session 1: mean difference= 0.01°/s, p=.92, BF_10_ = 2.4; Session 5: mean difference = 0.10°/s, t(9)=2.22, p=.054, BF_10_ =14.8).

### Adaptive responses are feature selective

A critical question raised by our findings is whether the learned compensatory response unveiled with a vertical Gabor (0°) is specifically triggered by the stimulus orientation feature, or whether it reflects a more general, unspecific motor response that can be driven by any moving target following sensorimotor training. To address this question, five of the same participants completed an additional control session where PRE and POST blocks included, in addition to the standard Gabor stimuli, trials with a Gaussian blob. The Gaussian blob is a stimulus with no orientation but otherwise identical motion properties. **Fig. 8** shows mean (across 5 participants) vertical eye velocity profiles for all four stimulus types in PRE and POST blocks within a single session. In the PRE block, all conditions showed vertical eye velocity near zero, apart from the tilted Gabors (45° and 135°) which exhibited the expected aperture-induced bias. Consistent with the 6 to 12-month delay since the last training session, the PRE block of Session 6 showed no residual compensatory response for the vertical Gabor, with vertical eye velocity remaining near zero across all non-tilted stimulus types, indicating that the learned motor command had not been retained over this extended timescale. In the POST training block, the vertical Gabor again showed a small but consistent upward shift in vertical eye velocity, replicating the compensatory response reported in **Fig. 7a**. More importantly, the vertical response to a moving blob remained indistinguishable from zero in both PRE and POST blocks, despite participants having undergone the same training with a +135° Gabor. Repeated measures ANOVA with Block (PRE, POST) and Orientation (0°, 45°, 135°, blob) as within-subject factors revealed a highly significant main effect of Orientation (F(3,12) = 26.29, *p_GG_* = .006), confirming robust stimulus-dependent differences in vertical eye velocity. The main effect of Block showed only a trend (F(1,4) = 5.34, p = .082), and the Block × Orientation interaction did not reach significance (F(3,12) = 2.21, *p_GG_* = .18). Post-hoc Tukey-corrected comparisons confirmed that 45° and 135° differed significantly from both 0° and blob (all p < .035), while 0° and blob did not significantly differ from each other (p = .053). Given the borderline nature of this comparison and our directional a priori hypothesis, we further examined this contrast with a planned paired t-test, which revealed a significant difference between POST 0° and POST blob (t(4) = 2.91, p = .044, Cohen’s d = 1.30). To complement the frequentist analyses, we computed Bayes factors using default Cauchy priors (r = 0.707). One-sample tests against zero indicated moderate evidence for a non-zero positive response in POST 0° (BF₁₀ = 3.99), while POST blob remained consistent with the null (BF₀₁ = 1.89). A paired Bayesian t-test directly comparing POST 0° to POST blob further supported a difference (BF₁₀ = 2.45). The absence of any POST response for the blob, contrasting with the positive shift observed for the vertical Gabor, demonstrates that the learned compensatory motor command is not triggered by motion alone but requires the presence of an orientation feature.

**Figure 8.**
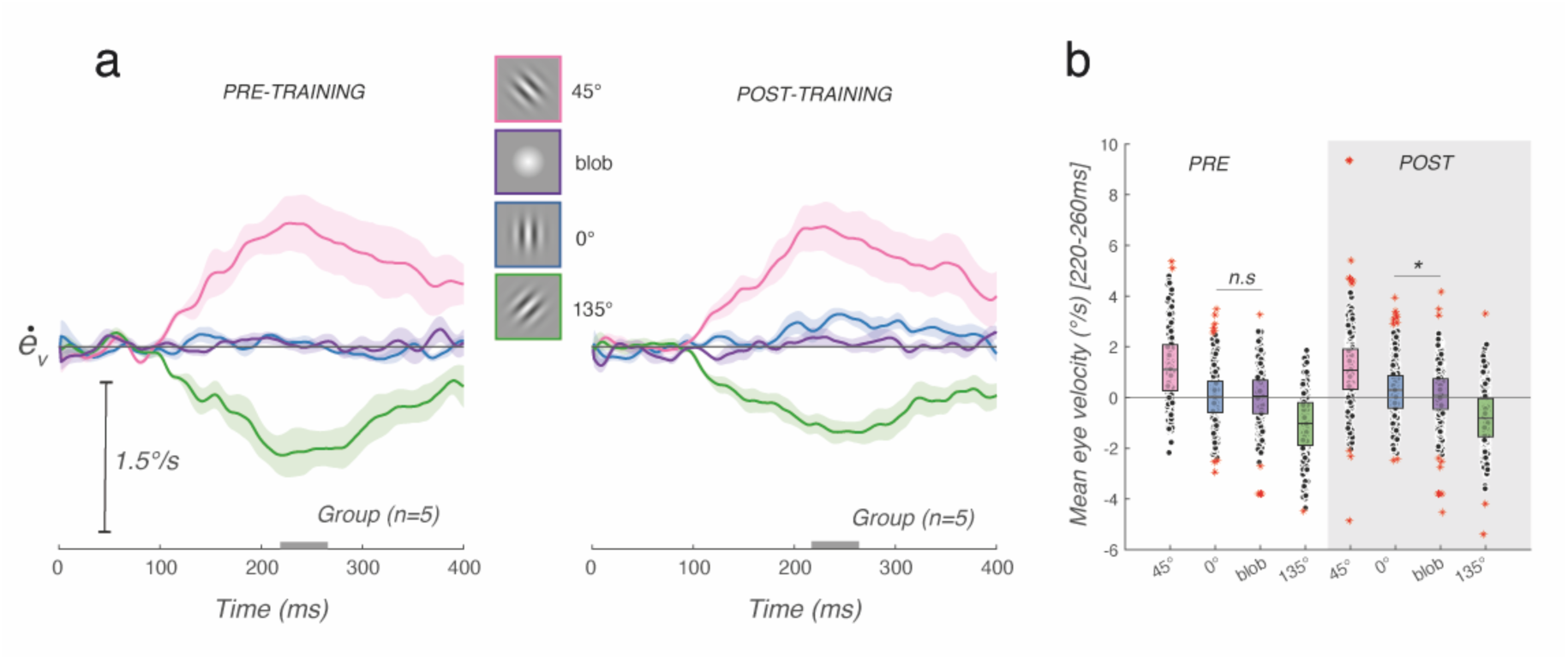
The learned compensatory response is stimulus-specific and requires orientation information. (**a**) Mean vertical eye velocity profiles for all four stimulus conditions: 45° Gabor (pink), 135° Gabor (green), vertical Gabor 0° (blue), and Gaussian blob (purple) averaged across 5 participants (n=5 for the blob session), for PRE and POST blocks of the additional Session 6. Shaded areas represent 85% confidence intervals. (**b**) Mean vertical eye velocity in the [220,260] msec window (closed-loop phase) for all conditions and both blocks, for all subjects.

### The compensatory response is replicated in an independent cohort

A remaining question is whether the compensatory response we describe generalizes beyond our original sample. To address this, we tested an independent group of 10 naïve participants, using a single session identical in structure to Session 1 of the main protocol (**Fig. 9**). As in the main cohort, tilted Gabors elicited the expected aperture-induced bias after pooling all the trials to the right, upward for 45° and downwards for 135°, while the vertical Gabor produced no measurable vertical eye velocity in the PRE block (**Fig. 9a**). Following a single training block of 720 trials with the 135° Gabor, a clear upward compensatory response emerged for the 0° Gabor (**Fig. 9a**, lower row), visible both in the [220,260] msec window measurements (**Fig. 9b**) and in the single-trial distribution with a shift towards the positive values after training (**Fig. 9c**). Mean vertical eye velocity for the vertical Gabor in the [220,260] msec window was significantly greater in POST than in PRE blocks (t(9) = 3.70, p=.005, BF_10_= 15.56). This replication in an independent sample demonstrates that the compensatory response is not idiosyncratic to our original participants, and that it can be induced within a single training session.

**Figure 9.**
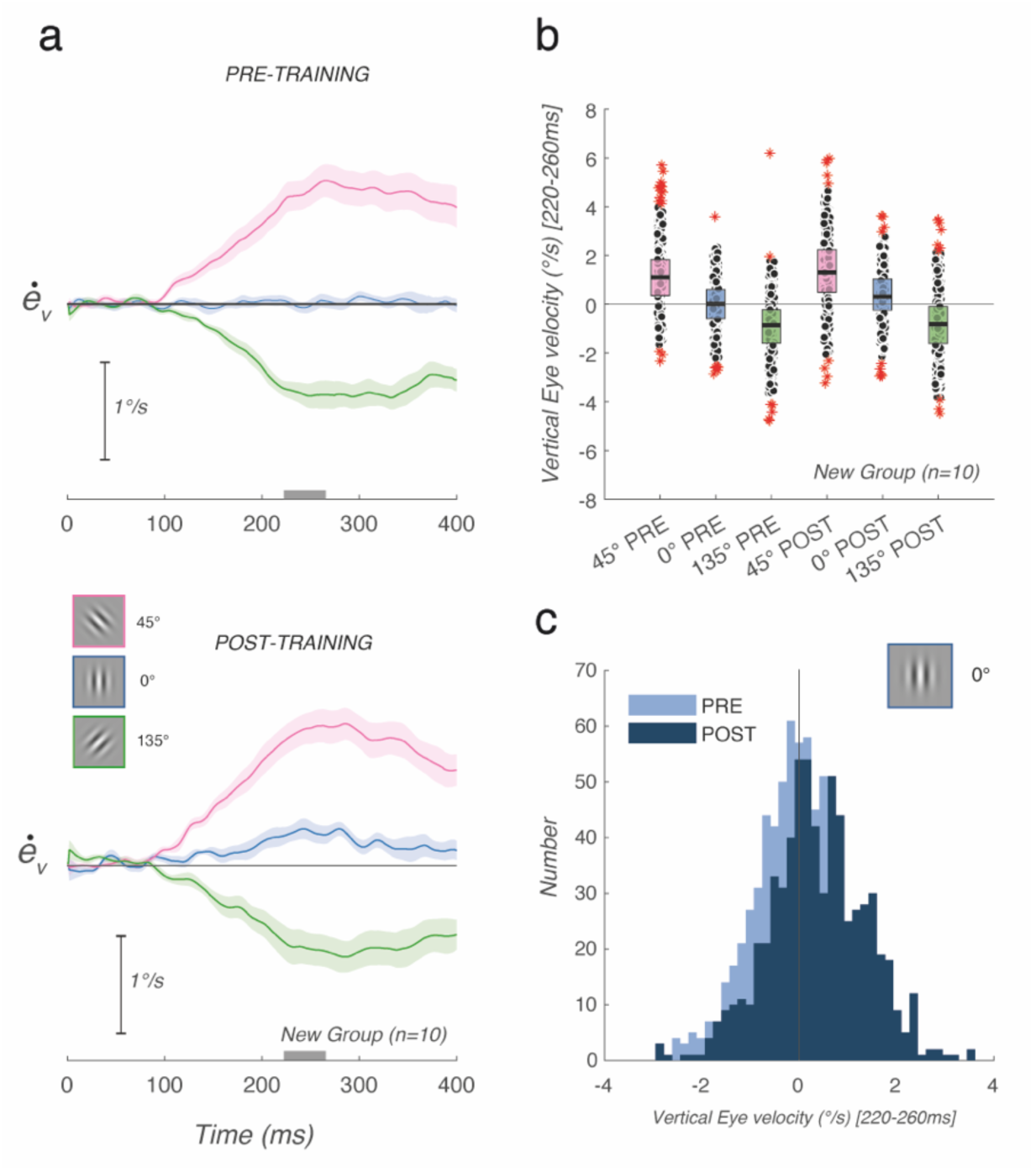
The compensatory response replicates in an independent cohort of naïve participants. **(a)** Mean vertical eye velocity profiles for the PRE and POST training blocks of a single session (S1), for all three Gabor orientations (blue: 0°, pink: 45°, green: 135°), averaged across 10 naïve participants. Shaded areas represent ±SEM. (**b**) Mean vertical eye velocity in the [220,260] msec window for all orientations and both blocks, for all participants. (**c)** Distribution of vertical eye velocity in the same window for the vertical Gabor, PRE (light blue) and POST (dark blue), pooled across all participants.

## DISCUSSION

Using Gabor stimuli, we confirmed that smooth pursuit is initially biased toward the direction orthogonal to target orientation, a hallmark of the aperture problem in motion perception ^11–13,28^. This bias reflects the tuning of early motion detectors, whose responses jointly encode orientation and direction ^14,29^. It is gradually resolved, such that within ∼300 msec pursuit direction closely matches true 2D target motion ^24,30^, mirroring the temporal dynamics of motion integration in macaque area MT ^14,20^. Within a Bayesian framework, this early misestimation is explained by combining local 1D motion signals with a slow-speed prior ^17^. Subsequent work has further shown that recurrent inference over temporally staggered local motion cues can account for both the dynamics of aperture-problem resolution and its behavioural consequences ^18,23^.

### Initial biases in motion direction computation cannot be cancelled

Crucially, the direction bias at pursuit onset proved resistant to extensive training: repeated exposure to a single orientation over hundreds of trials and multiple daily sessions left the initial error unchanged for both trained and untrained Gabor orientations. This finding contrasts with perceptual reports suggesting that motion-direction biases can be abolished by perceptual learning ^8,9^, although those studies largely relied on adjustment procedures, which may reflect shifts in decision criteria or motion adaptation rather than genuine changes in visual motion computations ^31^. By contrast, pursuit initiation during the open-loop interval provides a direct readout of early motion processing and its population decoding ^19,22,32^. Our results therefore argue that low-level sensory biases are not abolished by perceptual learning. Within a Bayesian framework, this would imply that sensory priors remain stable over hours to days despite sustained exposure to altered input statistics ^7^.

### But pursuit dynamics can still be adapted

That sensory biases resist cancellation does not mean that sensorimotor transformations are themselves immutable. The central finding of this study is that, although the initial directional bias at pursuit onset remained unchanged, the speed of its correction improved markedly with training, and did so selectively for the trained combination of orientation and motion direction. This improvement accumulated gradually across days, consistent with long-term sensorimotor learning rather than rapid perceptual recalibration. Within-session analyses further showed that residual direction error decreased progressively during training blocks, mirroring the across-session effect and suggesting a common error-correction mechanism across timescales. Notably, faster correction was not accompanied by improved final accuracy, which was already close to asymptote before training. Learning therefore altered the dynamics of compensation, not the sensory estimate itself.

The mechanism underlying this learning effect became apparent in responses to the vertical Gabor. Although a vertically oriented Gabor moving horizontally does not itself generate an aperture-induced direction bias, training elicited a robust vertical eye-velocity component to this stimulus. Its sign was opposite to the bias induced by the trained oblique orientation and generalized symmetrically across leftward and rightward motion. Repeated exposure to a stimulus that reliably drives an initial tracking error therefore builds a compensatory response in the opposite direction, accelerating error correction over time. In our paradigm, this adaptation was feature selective: it depended on both stimulus orientation and motion direction and was not elicited by a Gaussian blob lacking orientation information. This specificity argues against a simple low-level adaptation to repeated 1D motion signals ^31^ and instead supports a learned sensorimotor compensation for a predictable tracking error. To our knowledge, this is the first demonstration that smooth pursuit can acquire a compensatory response tailored to a systematic sensory bias.

These compensatory responses lagged the visually driven pursuit response by ∼50 ms. This delay is too short to be explained by visual feedback of the tracking error, which would require an erroneous eye movement, the generation of a retinal error signal, and subsequent correction through the visuomotor loop, a sequence expected to take at least 80–100 msec ^33,34^. Their timing and feature selectivity instead indicate that the compensatory command is triggered by the biased motor output itself. We therefore propose that learning establishes an internal model of the systematic motor error generated by the irreducible geometric ambiguity of the aperture problem. Because this ambiguity reflects an intrinsic property of local motion signals rather than uncertainty about the global target trajectory, it persists regardless of stimulus familiarity or training duration. The compensatory model, by contrast, builds gradually across days of training, consistent with our previous observation that learned expectations about 2D target trajectories can guide anticipatory pursuit without eliminating sensory misestimation caused by the aperture problem ^7^. Together, these results show that the pursuit system can learn both predictive and compensatory commands to optimize behaviour in the face of persistent sensory uncertainty. The control session performed months later further suggests that this learned association is not permanently retained but can be rapidly re-established within a single training block.

### Adaptive sensorimotor control: the hierarchical inference model

Our findings fit naturally within a hierarchical inference account of pursuit, in which initiation and steady-state tracking emerge from cascaded sensory and motor inferences shaped by prior knowledge, akin to Kalman filtering ^23,26^. Updating the slow-speed prior within recurrent sensory processing is sufficient to explain the gradual decay of tracking error during a trial ^18^. However, anticipatory pursuit before visual motion onset ^7,35–37^, as well as predictive responses to target disappearance and reappearance ^25,26^, point to an internal model learned from both prior responses and stimulus statistics. This model forms a probabilistic representation of the desired eye trajectory that matches the true 2D target path ^34,36^. It explains why, when target orientation and direction are predictable, anticipatory pursuit follows the veridical 2D trajectory even though subsequent visually driven responses remain biased by the aperture problem ^7^. Our results extend this framework by showing that the pursuit internal model can also learn to compensate for predictable distortions in sensory estimates, similar to other motor systems ^38^. The present training protocol should provide a useful model for probing the generalization principles of compensatory pursuit and object tracking through an implicit process ^39^ .

Our results also inform the neural circuitry underlying Bayesian-like pursuit behaviour. The frontal eye fields (FEF), and in particular its smooth-pursuit subregion FEF_SEM_, are strong candidates for expressing the learned compensatory command. FEF contributes to anticipatory pursuit initiation, dynamic gain control and velocity maintenance ^40,41^, and receives input from feature-selective visual areas that could provide the conditioned signals driving the learned association. A plausible circuit therefore comprises three interacting inference nodes: MT, implementing recurrent sensory inference ^14^; prefrontal and frontal eye-field networks, computing and updating desired trajectories ^42^; and cerebellar–pontine pathways, consolidating the association learning across sessions through error-driven plasticity ^32,43^. The present paradigm offers an experimental framework for disentangling their respective contributions.

## Conclusion

Whether perceptual illusions influence sensorimotor transformations has long been debated in the perception–action literature. Here we exploited the aperture problem, an intrinsic and generic feature of many sensory computations ^44^, to test how sensorimotor systems compensate for predictable yet unavoidable misestimation. Smooth pursuit emerges from the interplay of sensory and motor internal models that are flexibly weighted by context ^25,26,32^. Together, our findings show that these interacting computational stages implement an adaptive, biologically plausible form of Kalman filtering whose principles likely extend far beyond oculomotor control to other sensorimotor transformations ^45^.

## Acknowledgments

This work was first presented at the annual Vision Sciences Society meeting in May 2025 (Schoeffel et al., 2025). This research was funded by a joint NSF–ANR international grant from the CRCNS programme (PrioSens, ANR-20-NEUC-002-01) awarded to G.S.M.. C.L.S. was supported by a fellowship from the Fondation de France. A.M. was supported by the collaborative grant Vision-3E, ANR-21-CE37-0018-01. We thank Andrew Meso, Philippe Lefevre and Pascal Mamassian for their critical reading of the manuscript.

## MATERIALS AND METHODS

### Participants

10 participants (5 male, 5 female; mean age, 29 ± 10 years) took part in this study. All had normal or corrected-to-normal vision and no known neurological or ophthalmological disorders. The study was approved by an ethics committee (Priosens, CPP Nord-Ouest 1, 2022-A01586-37) under French regulations governing research involving human participants. All participants provided written informed consent in accordance with the Declaration of Helsinki and were compensated at a rate of €10 per hour. Notice that the identification using numbers S01 to S27 follows the naming of the available dataset, following the BIDS format.

### Display and apparatus

Visual stimuli were generated in MATLAB (The MathWorks Inc., Natick, MA) using Psychtoolbox-3 on a Mac computer. Stimuli were displayed on a calibrated DISPLAY++ LCD monitor (Cambridge Research Systems; resolution, 1920 × 1080 pixels; refresh rate, 85 Hz) at a viewing distance of 57.9 cm, subtending approximately 62° of visual angle. Experiments were conducted in a dark booth. Movements of the right eye were recorded with a tower-mounted EyeLink 1000 Plus video eye tracker (SR Research, Canada) at a sampling rate of 1000 Hz. Participants’ heads were stabilized with a chin-and-forehead rest, and stimuli were viewed through a hot mirror that also allowed imaging of the eye.

### Motion stimuli

Moving targets were Gabor patches consisting of sinusoidal gratings multiplied by a Gaussian envelope (diameter: ∼5° of visual angle, σ = 0.84°, spatial frequency: 0.3 cycles/°; Michelson contrast: 50%). Gabors were presented at three orientations: vertical (0°) and two oblique tilts (45° and 135°). Stimuli translated horizontally (leftwards and rightwards, equiprobable across trials) at a constant speed of 10°/s.

### Experimental design and procedure

Each trial began with a 200 msec central fixation period, during which participants fixated a small Gaussian blob (0.5° FWHM) displayed at the centre of the screen. The fixation stimulus then disappeared, followed by a 300 msec gap before the Gabor target appeared at the same location and began translating horizontally. This gap paradigm was used to facilitate smooth-pursuit initiation by disengaging fixation and reducing the likelihood of catch-up saccades at motion onset. Participants were instructed to track the moving target as smoothly and accurately as possible throughout each trial. Target motion lasted 500 msec, sufficient to reach steady-state tracking, after which the target disappeared and the fixation spot reappeared, signalling the start of the next trial (**Fig. 1a**).

Each session consisted of 1200 trials organized into three successive blocks: a pre-training block (240 trials), a training block (720 trials), and a post-training block (240 trials), lasting approximately one hour in total. Each participant completed 5 sessions (∼6000 trials total) over several days. The identical structure of the pre- and post-training blocks across sessions allowed estimation of learning effects both within and across sessions.

In the pre- and post-training blocks, all three Gabor orientations (0°, 45°,135°) and both motion directions were fully interleaved to prevent anticipatory pursuit and to provide an unbiased estimate of tracking performance across the full orientation set. Each block was divided into three sub-blocks of 80 trials each, with short breaks between sub-blocks and recalibration to prevent eye fatigue.

### Learning Paradigm

The training block was designed to induce perceptual learning for one specific Gabor orientation (**Fig 1b**). During training, a single oblique orientation (135°) was presented repeatedly over all 720 trials, with motion direction (leftward and rightward) randomized across trials. The training block was divided into nine sub-blocks of 80 trials each, with short breaks and recalibration between sub-blocks to prevent eye fatigue. Two separate series of 5 sessions were conducted in independent groups of 5 participants: in the first series, training speed matched the pre/post baseline (10°/s); in the second series, training speed was doubled (20°/s). This speed manipulation was designed to test two hypotheses: (i) that prolonged exposure to a single orientation of the moving target selectively reduces the direction bias associated with that orientation via perceptual learning, and (ii) that pairing a specific orientation with a higher speed during training may facilitate bias correction.

### Blob control session

A subset of 5 participants completed one additional session 6–12 months after completing the five main learning sessions. This session followed the same structure as the standard pre- and post-training blocks, with the exception that PRE and POST blocks included an additional stimulus type: a Gaussian blob (diameter: 4.5°; σ = 1.17°) moving horizontally at 10°/s, presented with equal probability alongside the standard Gabor stimuli. Each PRE and POST block therefore contained 40 additional trials per direction, for a total of 80 additional trials per block. The blob stimulus had no oriented carrier and thus carried no aperture-induced directional ambiguity but was otherwise identical to the Gabor stimuli in terms of motion speed, direction, and duration. The extended delay between the main protocol and this control session ensured that any long-term retention of the learned compensatory response had dissipated, allowing us to test whether a single training block could rapidly reinduce the orientation-specific compensatory response.

### Replication cohort

An independent group of 10 naïve participants (4 male, 6 female; mean age, 29 ± 6 years) completed a single session identical in structure to Session 1 of the main protocol (PRE block, 240 trials; training block at 10°/s, 720 trials with the 135° Gabor; POST block, 240 trials), to test whether the compensatory response reported above generalizes to an independent sample.

### Movement analysis

*Preprocessing.* Horizontal and vertical eye position signals were low-pass filtered using a second-order Butterworth filter (cutoff: 25 Hz) applied via zero-phase filtering. Filter quality was verified by computing, for each trial, the Pearson correlation between raw and filtered position traces and the root-mean-square (RMS) of the residual (raw – filtered): mean correlations were r = 0.9997 (horizontal) and r = 0.9903 (vertical), with mean RMS residuals of 0.05° and 0.02° respectively, confirming that the filter removed high-frequency tracker noise without distorting eye movement dynamics. Eye velocity was estimated using a four-point central difference method ^46^: for each time point *k*, velocity was computed as :

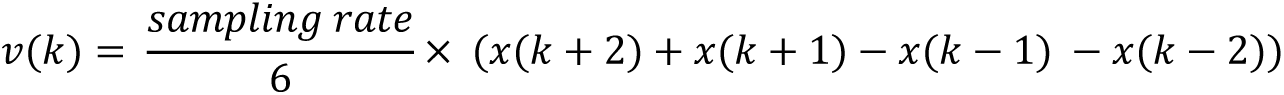

This method provides a smooth and stable instantaneous velocity estimate at 1 kHz while preserving temporal resolution. Missing data due to blinks or signal dropouts propagated as NaN values into the velocity trace, ensuring that corrupted segments were automatically excluded from further analysis. Saccades were detected using a velocity threshold criterion (30°/s) and replaced by linear interpolation.

#### Pursuit velocity and direction

Pursuit velocity was computed as the mean vertical eye velocity within 12 successive 40 ms time windows, with onsets ranging from 0 to 480 msec after motion onset. Initial pursuit velocity and direction were taken from the 140 to 180 msec post-onset window, which falls in the open-loop, accelerating phase of the smooth pursuit given a mean latency of ∼100 msec. Tracking direction was computed per trial and per time window. Because Gabor orientation had no significant effect on pursuit latency, velocity and direction estimates were computed relative to motion onset consistently across all orientation conditions. Latency estimates were also extracted manually for the vertical component of eye velocity separately for each orientation, block, and session, and compared across conditions to characterize the onset timing of both the aperture-induced bias and the learned compensatory response.

*Exponential fitting of correction dynamics*

To characterize the temporal dynamics of direction error correction, the time course of mean eye velocity was fitted with an exponential function of the form:

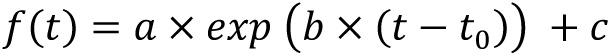

Where *a* is the initial error amplitude, *b* the decay rate (from which the time constant 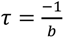 is derived), *c* the residual (steady state) error, and *t*_0_ the time origin of the fit. Fits were performed on mean tracking angle computed in successive 40 msec time bins spanning from 120 to 440 msec after motion onset, within initial parameter estimates determined by identifying the best-fitting values within the (120-240) ms windows. This model was applied separately for each orientation (45° and 135°) and direction condition, for each participant. Fits were inspected individually and excluded from further analyses when they failed to converge, produced implausible parameter values, or captured the wrong direction change. This yielded n=7 valid fits per condition across subjects.

*Gamma fitting of pursuit velocity profiles*

To characterize the full temporal profile of mean eye velocity across all orientation conditions, a Gamma function was fitted to the eye velocity time course:

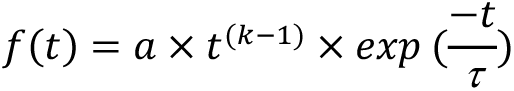

Where (*a*) is a scaling factor, (*k*) the shape parameter controlling the rise time, and (τ) the scale parameter controlling the decay. Fits were performed on mean eye velocity computed in successive 40 ms time bins spanning from 40 to 480 msec after motion onset, covering the full pursuit response from initiation through steady state. Unlike the exponential model applied to direction error correction, this model captures the full rise-and-fall dynamics of the pursuit velocity response and was applied across all orientations (0°, 45°, 135°).

### Quantification and Statistical Analysis

All statistical analyses were performed in MATLAB. The alpha threshold was set at 0.05. Where multiple comparisons were performed, p-values were corrected using Holm-Bonferroni correction unless otherwise stated. Repeated-measures ANOVAs were corrected for sphericity violations using the Greenhouse-Geisser method where applicable; corrected p-values are reported as *p_GG_* throughout.

To quantify evidence for both significant and null results, Bayes factors (BF_10_) were computed for all statistical tests using a Bayesian Information Criterion (BIC) approximation method implemented in MATLAB. For ANOVA designs, Bayes factors were derived from the sums of squares and degrees of freedom associated with each effect relative to the error term. For linear mixed-effects models, BF_10_ values were estimated by comparing the BIC of the full model with reduced models excluding the factor of interest. In all cases, BF_10_ represents the evidence in favour of the alternative hypothesis relative to the null hypothesis. Evidence strength was interpreted using conventional thresholds (Jeffreys, 1961; Kass & Raftery, 1995), with BF_10_> 3 indicating positive evidence for the alternative hypothesis and BF_10_ < 0.33 indicating positive evidence for the null hypothesis.

#### Baseline pursuit dynamics (***Fig 2***)

To characterize the directional bias induced by Gabor orientation, vertical eye velocity (mean over the 180-220ms window) was analysed using two-way repeated-measures ANOVAs with Orientation (0°, 45°, 135°) and Direction (Left, Right) as within-subject factors (n=10). The [180,220] msec window was chosen as it falls at the end of the open-loop phase, where the aperture-induced vertical eye velocity has reached its peak amplitude before the closed-loop correction begins, maximizing sensitivity to the direction bias. The [140,180] msec window used for learning analyses captures the earlier, accelerating phase of pursuit initiation. The same design was applied to the parameters of exponential fits to the velocity time course: initial error (*a*), residual error (*c*), and log-transformed time constant (τ), restricted to the two oblique orientations (45°, 135°; n= 6-7 subjects after exclusion of invalid fits). Significant interactions were followed up with paired t-tests, Holm corrected. Single-subject analyses used trial-level two-way factorial ANOVAs with the same factors (∼ 50 trials per condition).

#### Between-group comparison (20°/s vs 10°/s)

To compare participants trained at high speed (20°/s) versus normal speed (10°/s), a linear mixed-effects model was used with Group as a between-subject factor and Session and Block as within-subject factors, with random intercepts per subject. Dependent variables (mean eye velocity, initial tracking error *a*, and time constant τ) were tested in separate models.

#### Learning effects (***Fig 3***)

To assess the effects of perceptual learning on pursuit, mean eye velocity in the [140,180] msec window (i.e. a 40msec long time window) was analysed using three-way repeated-measures ANOVAs with Block (PRE, POST), Session (S1, S5), and Orientation (0°, 45°, 135°) as within-subject factors (n=10). Significant higher-order interactions were decomposed using simple effects (paired t-tests, Bonferroni corrected). For the vertical orientation (0°) specifically, a 2 x 2 repeated-measures ANOVA (Session x Block) was conducted at the group level (n=10) to assess the temporal evolution of the compensatory response. Single-subject analyses used trial-level factorial ANOVAs with the same factor structure. As a model-free complement to the exponential decay analysis, we computed t_50%_, the time at which the pursuit angle reached the midpoint between its initial value and its asymptote. We analysed it using a three-way repeated-measures ANOVA with Session (Session 1-5), Phase (PRE, POST) and Orientation (45°, 135°) as within-subject factors.

#### Time constant reliability (***Fig 4***)

To assess whether individual convergence dynamics are stable across learning, Pearson correlations between PRE and POST time constants (τ) were computed separately for each oblique orientation (45°, 135°), at both the single-subject and group level.

#### Linear superposition analyses (***Fig 7*)**

We tested whether the post-learning velocity response to the trained orientation (POST 135°) could be predicted by a linear combination of the response to the vertical control (POST 0°) and the pre-learning response to the trained orientation (PRE 135°). Pearson correlations between predicted and observed values were computed both at the single-subject and group level (pooled across subjects).

#### Vertical eye velocity latency analysis (***Fig 7***)

Pursuit onset latencies for the vertical eye velocity component were estimated manually from mean velocity profiles for each participant, session, block, and orientation. Latency differences between the vertical Gabor (0°) and oblique orientations (45°, 135°) were assessed using paired t-tests on subject means, with Bayes factors computed using a JZS prior (r^2^ = 0.707). A repeated-measures ANOVA was used to test for effects of Orientation and Session on latency, with post-hoc comparisons using multiple comparison tests.

#### Blob control session (***Fig. 8***)

To test whether the learned compensatory response is specific to oriented stimuli, mean vertical eye velocity in the [220,260] msec window was analysed using a two-way repeated-measures ANOVA with Block (PRE, POST) and Orientation (0°, 45°, 135°, blob) as within-subject factors (n=5). Normality was assessed using Lilliefors tests for each condition. Post-hoc pairwise comparisons between orientations were conducted using the MATLAB multiple comparison tests with Bonferroni correction.

#### Replication cohort (***Fig. 9***)

To assess whether the compensatory response generalizes to an independent sample, mean vertical eye velocity in the [220,260] msec window was compared between PRE and POST blocks for the 0° Gabor, with leftward trials sign-flipped and pooled with rightward trials. For each participant, a single mean value was computed per block, and PRE and POST values were compared across the 10 participants using a paired t-test. Normality of the paired differences was verified using a Lilliefors test. Bayes factors were computed using a JZS prior (r = 0.707).

## Notes

### Competing Interest Statement

The authors have declared no competing interest.

### Summary of Updates

A new statistical analysis (single subject level) has been performed on the original data set, confirming the results. Moreover, a replication study involving a new group of 10 naive participants has been ran, confirming the original results that a delayed, compensatory pursuit eye movement emerge after the first training session.

## REFERENCES

1. Born, R. T. C, Bencomo, G. M. Illusions, Delusions, and Your Backwards Bayesian Brain: A Biased Visual Perspective. Brain. Behav. Evol. 95, 272–285 (2020).

2. Gregory, R. L. Visual illusions classified. Trends Cogn. Sci. 1, 190–194 (1997).

3. Mamassian, P., Landy, M. C, Maloney, L. T. Bayesian Modelling of Visual Perception. in Probabilistic Models of the Brain (eds Rao, R. P. N., Olshausen, B. A. C, Lewicki, M. S.) 13–36 (The MIT Press, 2002). doi:10.7551/mitpress/5583.003.0005.

4. Körding, K. P. C, Wolpert, D. M. Bayesian decision theory in sensorimotor control. Trends Cogn. Sci. 10, 319–326 (2006).

5. Milner, D. C, Dyde, R. Why do some perceptual illusions affect visually guided action, when others don’t? Trends Cogn. Sci. 7, 10–11 (2003).

6. Chalk, M., Seitz, A. R. C, Series, P. Rapidly learned stimulus expectations alter perception of motion. J. Vis. 10, 2–2 (2010).

7. Montagnini, A., Spering M. C, Masson, G. S. Predicting 2D Target Velocity Cannot Help 2D Motion Integration for Smooth Pursuit Initiation. J. Neurophysiol. 96, 3545– 3550 (2006).

8. Seriès, P. C, Seitz, A. R. Learning what to expect (in visual perception). Front. Hum. Neurosci. 7, (2013).

9. Sotiropoulos, G., Seitz, A. R. C, Seriès, P. Changing expectations about speed alters perceived motion direction. Curr. Biol. 21, R883–R884 (2011).

10. Szpiro, S. F. A., Spering, M. C, Carrasco, M. Perceptual learning modifies untrained pursuit eye movements. J. Vis. 14, 8–8 (2014).

11. Wallach, H. Über visuell wahrgenommene Bewegungsrichtung. Psychol. Forsch. 20, 325–380 (1935).

12. Adelson, E. H. C, Movshon, J. A. Phenomenal coherence of moving visual patterns. Nature 300, 523–525 (1982).

13. Castet, E., Charton, V. C, Dufour, A. The extrinsic/intrinsic classification of two-dimensional motion signals with barber-pole stimuli. Vision Res. 39, 915–932 (1999).

14. Pack, C. C. C, Born, R. T. Temporal dynamics of a neural solution to the aperture problem in visual area MT of macaque brain. Nature 409, 1040–1042 (2001).

15. Tlapale, É., Masson, G. S. C, Kornprobst, P. Modelling the dynamics of motion integration with a new luminance-gated diffusion mechanism. Vision Res. 50, 1676– 1692 (2010).

16. Stocker, A. A. C Simoncelli, E. P. Noise characteristics and prior expectations in human visual speed perception. Nat. Neurosci. 9, 578–585 (2006).

17. Weiss, Y., Simoncelli, E. P. C, Adelson, E. H. Motion illusions as optimal percepts. Nat. Neurosci. 5, 598–604 (2002).

18. Montagnini, A., Mamassian, P., Perrinet, L., Castet, E. C, Masson, G. S. Bayesian modeling of dynamic motion integration. J. Physiol.-Paris 101, 64–77 (2007).

19. Perrinet, L. U. C Masson, G. S. Motion-Based Prediction Is Sufficient to Solve the Aperture Problem. Neural Comput. 24, 2726–2750 (2012).

20. Smith, M. A., Majaj, N. J. C, Movshon, J. A. Dynamics of motion signaling by neurons in macaque area MT. Nat. Neurosci. 8, 220–228 (2005).

21. Solomon, S. S. et al. Visual motion integration by neurons in the middle temporal area of a New World monkey, the marmoset. J. Physiol. 589, 5741–5758 (2011).

22. Barthélemy, F. V., Perrinet, L. U., Castet, E. C, Masson, G. S. Dynamics of distributed 1D and 2D motion representations for short-latency ocular following. Vision Res. 48, 501–522 (2008).

23. Bogadhi, A. R., Montagnini, A., Mamassian, P., Perrinet, L. U. C, Masson, G. S. Pursuing motion illusions: a realistic oculomotor framework for Bayesian inference. Vision Res. 51, 867–880 (2011).

24. Masson, G. S. C Stone, L. S. From Following Edges to Pursuing Objects. J. Neurophysiol. 88, 2869–2873 (2002).

25. Bogadhi, A. R., Montagnini, A. C, Masson, G. S. Dynamic interaction between retinal and extraretinal signals in motion integration for smooth pursuit. J. Vis. 13, 5–5 (2013).

26. Orban de Xivry, J.-J., Coppe, S., Blohm, G. C, Lefèvre, P. Kalman Filtering Naturally Accounts for Visually Guided and Predictive Smooth Pursuit Dynamics. J. Neurosci. 33, 17301–17313 (2013).

27. Patricio Décima, A., Fernando Barraza, J. C López-Moliner, J. The perceptual dynamics of the contrast induced speed bias. Vision Res. 191, 107966 (2022).

28. Lorenceau, J., Shiffrar, M., Wells, N. C, Castet, E. Different motion sensitive units are involved in recovering the direction of moving lines. Vision Res. 33, 1207–1217 (1993).

29. Albright, T. D. Direction and orientation selectivity of neurons in visual area MT of the macaque. J. Neurophysiol. 52, 1106–1130 (1984).

30. Wallace, J. M., Stone, L. S. C, Masson, G. S. Object Motion Computation for the Initiation of Smooth Pursuit Eye Movements in Humans. J. Neurophysiol. 93, 2279– 2293 (2005).

31. Gekas, N. C Mamassian, P. Adaptation to one perceived motion direction can generate multiple velocity aftereffects. J. Vis. 21, 17 (2021).

32. Lisberger, S. G. Toward a Biomimetic Neural Circuit Model of Sensory-Motor Processing. Neural Comput. 35, 384–412 (2023).

33. 33. Goldreich, D., Krauzlis, R. J. C, Lisberger, S. G. Effect of changing feedback delay on spontaneous oscillations in smooth pursuit eye movements of monkeys. (1992).

34. Perrinet, L. U., Adams, R. A. C, Friston, K. J. Active inference, eye movements and oculomotor delays. Biol. Cybern. 108, 777–801 (2014).

35. Carneiro Morita, V., Souto, D., Masson, G. S. C, Montagnini, A. Anticipatory smooth pursuit eye movements scale with the probability of visual motion: The role of target speed and acceleration. J. Vis. 25, 2 (2025).

36. Damasse, J.-B., Perrinet, L. U., Madelain, L. C, Montagnini, A. Reinforcement effects in anticipatory smooth eye movements. J. Vis. 18, 14 (2018).

37. Pasturel, C., Montagnini, A. C, Perrinet, L. U. Humans adapt their anticipatory eye movements to the volatility of visual motion properties. PLOS Comput. Biol. 16, e1007438 (2020).

38. Shadmehr, R., Smith, M. A. C, Krakauer, J. W. Error Correction, Sensory Prediction, and Adaptation in Motor Control. Annu. Rev. Neurosci. 33, 89–108 (2010).

39. Wilterson, S. A. C, Taylor, J. A. Implicit Visuomotor Adaptation Remains Limited after Several Days of Training. eneuro 8, ENEURO.0312-20.2021 (2021).

40. MacAvoy, M. G., Gottlieb, J. P. C, Bruce, C. J. Smooth-Pursuit Eye Movement Representation in the Primate Frontal Eye Field. Cereb. Cortex 1, 95–102 (1991).

41. Nuding, U. et al. TMS Evidence for Smooth Pursuit Gain Control by the Frontal Eye Fields. Cereb. Cortex 19, 1144–1150 (2009).

42. Yang, J., Lee, J. C, Lisberger, S. G. The Interaction of Bayesian Priors and Sensory Data and Its Neural Circuit Implementation in Visually Guided Movement. J. Neurosci. 32, 17632–17645 (2012).

43. Kahlon, M. C, Lisberger, S. G. Changes in the Responses of Purkinje Cells in the Floccular Complex of Monkeys After Motor Learning in Smooth Pursuit Eye Movements. J. Neurophysiol. 84, 2945–2960 (2000).

44. Pack, C. C. C, Bensmaia, S. J. Seeing and Feeling Motion: Canonical Computations in Vision and Touch. PLOS Biol. 13, e1002271 (2015).

45. Wolpert, D. M. C, Flanagan, J. R. Computations underlying sensorimotor learning. Curr. Opin. Neurobiol. 37, 7–11 (2016).

46. Engbert, R. C, Kliegl, R. Microsaccades uncover the orientation of covert attention. Vision Res. 43, 1035–1045 (2003).

